# Polygenic Risk Scores for Kidney Function to the Circulating Proteome, and Incident Kidney Diseases: the Atherosclerosis Risk in Communities Study

**DOI:** 10.1101/2020.09.05.284265

**Authors:** Zhi Yu, Jin Jin, Adrienne Tin, Anna Köttgen, Bing Yu, Jingsha Chen, Aditya Surapaneni, Linda Zhou, Christie M. Ballantyne, Ron C. Hoogeveen, Dan E. Arking, Nilanjan Chatterjee, Morgan E. Grams, Josef Coresh

## Abstract

Genome-wide association studies (GWAS) have revealed numerous loci for kidney function (estimated glomerular filtration rate, eGFR). The relationship of polygenic predictors of eGFR, risk of incident adverse kidney outcomes, and the plasma proteome is not known. We developed a genome-wide polygenic risk score (PRS) using a weighted average of 1.2 million SNPs for eGFR using the LDpred algorithm, summary statistics generated by a European-ancestry (EA) meta-analysis of the CKDGen Consortium (N=558,423) and UK Biobank GWAS for eGFR (90% of the cohort; N=289,432), followed by best parameter selection using data from the remaining 10% of the UK Biobank (N=32,159). We then tested the association of the PRS among 8,886 EA participants in the Atherosclerosis Risk in Communities (ARIC) study (mean age: 54±6 years, 53% female) with incident chronic kidney disease (CKD), end stage kidney disease (ESKD), kidney failure (KF), and acute kidney injury (AKI). We also examined 4,877 plasma proteins measured at two time points (visit 3 (1993-95) and visit 5 (2011-13)) in relation to the PRS and compared associations between the proteome and eGFR itself. All models were adjusted for age, sex, center, and the first 10 principal components of ancestry. The developed PRS had an R^2^ for eGFR of 0.07 in ARIC. Over 30 years of follow up, the number of incident CKD, ESKD, KF, and AKI were 2,959, 137, 470, and 1,723, respectively. The PRS showed significant associations with all outcomes: hazard ratios (95% CI) per 1 SD lower PRS were 1.33 (1.28, 1.39), 1.20 (1.00, 1.42), 1.17 (1.06, 1.28), and 1.07 (1.02, 1.12) for incident CKD, ESKD, KF, and AKI respectively. The PRS was significantly associated (Bonferroni threshold P<1.02 × 10^−5^) with 108 proteins at both time points. The strongest associations were with cystatin-C (a marker of kidney function used in clinical practice), collagen alpha-1 (XV) chain, and desmocollin-2. All significant correlations with the PRS were negative, except those of testican-2 and angiostatin. Correlations of proteins with eGFR were much stronger than those with the PRS. Overall, we demonstrated that the PRS for eGFR is now sufficiently strong to capture risk for a spectrum of incident kidney diseases as well as broadly influence the plasma proteome.

## INTRODUCTION

Most kidney diseases are complex diseases with both genetic and environmental factors contributing to their risks. The heritability estimated by family studies are 30–75%.^1-4^ Genomewide associations studies (GWAS) have grown rapidly in the last decade and identified numerous loci for kidney function, which gave rise to increasing attention to testing polygenic risk scores (PRS) as risk factors for kidney diseases risks.^5-11^ However, previous PRS provided limited risk stratification for adverse kidney outcomes such as end-stage kidney disease (ESKD).^5,10^ Potential reasons include: small sample sizes of early GWAS, which might lead to imprecise estimation of the associations between individual variants and disease risk; limiting the PRS to genetic variants that reached genome-wide significance (P < 5 × 10^−8^); and a lack of deeply phenotyped data to identify cases.^6-11^ With new data and methodologies, there is an opportunity to mitigate these limitations.

New methodologies for large scale proteomic measurement using aptamer technologies also provide an opportunity to assess the impact of genetic susceptibility to low kidney function on the plasma proteome.^12,13^ The plasma proteome consists of thousands of secreted proteins that involve in numerous physiological and pathological processes, including transporting and signaling, metabolism, vascular function, and defense mechanisms.^14-16^ Therefore, the plasma proteome is a reservoir of important potential biomarkers capturing current physiology and pathophysiology. Although previous studies have demonstrated the heritability of plasma protein levels,^17^ research into the plasma proteomic signals of genetic susceptibility including that for kidney diseases, has been limited.^18-21^ In kidney diseases, as reduced kidney function is correlated with elevations in many proteins but the balance of genetically predicted risk vs. secondary influences on proteomic signals is unknown.

Using large studies and new algorithms, we investigated the strength of associations of PRS for kidney function with incident kidney diseases over 30 years of follow-up in a deeply phenotyped community-based cohort. We included diseases strongly related to kidney function with evidence of a strong genetic basis, including chronic kidney disease (CKD), end-stage kidney disease (ESKD), and kidney failure,^1-4^ as well as acute kidney injury (AKI).^22^ We also examined 4,877 plasma proteins measured at two time points approximately 20 years apart in relation to both genetic susceptibility to low kidney function and the concurrent kidney function itself, in order to evaluate the strengths of proteomic associations with genetically predicted risk and physiological changes and how those associations change over time.

## METHODS

### Study Cohort

The Atherosclerosis Risk in Communities (ARIC) study is an ongoing longitudinal cohort of 15,792 45-60-year-old participants (55% female, 73% participants of European ancestry (EA)) recruited from four communities in the U.S.: Forsyth County, North Carolina; Jackson, Mississippi; suburbs of Minneapolis, Minnesota; and Washington County, Maryland at 1987-1989 (visit 1). Follow-up examinations were conducted approximately every three years: 1990-1992 (visit 2), 1993-1995 (visit 3), 1996-1998 (visit 4), more recently, in 2011-2013 (visit 5), in 2016-2017 (visit 6), and in 2018-2019 (visit 7).^23^ Each study visit consisted of a clinical examination, blood and urine specimens collection, and filling out extensive questionnaires. Proteomic levels were measured at visit 3 and visit 5. Our primary study population was restricted to 8,866 unrelated EA participants (**Supplemental Figure 1**), since so far most of the genomic studies with results available for use were conducted among EA. In the proteomic analysis of our study, 7,213 participants with valid proteomic measurements remained. In sensitivity analysis, we constructed another study population of 2,871 unrelated participants with African ancestry (AA). Study protocols were approved by the Institutional Review Boards and all study participants provided informed consent (including agreement for industry studies for SomaLogic sponsored proteomic quantification).

### Genotyping

Participants were genotyped with the Affymetrix 6.0 DNA microarray (Affymetrix, Santa Clara, CA) with genotype calling performed using the Birdseed algorithm. Genotyping was performed on the Affymetrix 6.0 DNA microarray (Affymetrix, Santa Clara, CA) and analyzed with the Birdseed variant calling algorithm. Haplotype phasing was performed using ShapeIt (v1.r532).^24^ Genotypes were imputed on the Michigan Server to the TOPMed reference panel.^25,26^ A quality control was carried out prior to imputation: SNPs were included if they had call rate < 95%, Hardy-Weinberg equilibrium p-values < 0.0001, or minor allele frequencies (MAF) < 1%.^27^ Individuals with cryptic relatedness defined as identity by state (IBS) distance generated from PLINK > 0.8 were also excluded.^28^

### Assessing Kidney Function

Kidney function, measured as estimated glomerular filtration rate (eGFR), was assessed by measuring serum creatinine (at all visits excepted visit 7) and serum cystatin C (at all visits excepted visit 1 and 7) using the 2009 Chronic Kidney Disease Epidemiology Collaboration (CKD-EPI) creatinine equation (eGFRcr) and 2012 CKD-EPI cystatin C equation (eGFRcys).^29,30^ Cystatin C is an excellent marker for kidney function but is not as widely used as creatinine, which limits its use in genetics studies. Serum creatinine level was measured by the modified kinetic Jaffé method, standardized to the National Institute of Standards and Technology (NIST) standard, and calibrated to an isotope dilution mass spectrometry (IDMS)-traceable reference method.^31-33^ Serum cystatin C level was measured by the turbidimetric method, and standardized and calibrated to the International Federation of Clinical Chemistry and Laboratory Medicine (IFCC) reference.^34^ In the polygenic risk scores development, we used eGFRcr as the kidney function measurement since this has been the main trait with the largest samples size in GWAS meta-analysis.^5^

### Polygenic Risk Score for Kidney Function

Polygenic risk scores aggregate genome-wide genetic variation into a single score that reflects individual’s inherited disease risk. They are most commonly calculated by summing across SNPs associated with a given trait, weighted by their effect sizes from GWAS results of that trait.

For the PRS construction, we first conducted a GWAS for log(eGFRcr) using PLINK among 90% of unrelated EA participants in the UK Biobank (N=289,432; application ID 17712) using an additive genetic model adjusted for age and sex.^28^ Details of the UK Biobank cohort has been described elsewhere.^35^ Then we conducted a fixed-effects inverse variance weighted metaanalysis on the summary statistics from the UK Biobank GWAS and a meta-analysis by the CKDGen Consortium of the GWAS of eGFRcr including up to 567,460 EA individuals.^5,36^ As the CKDGen consortium included the ARIC study (N=9,037), we adjusted the effect sizes of SNPs by removing the ARIC participants.^37^ We also used a sample of 489 unrelated EA individuals from phase 3 1000 Genomes as a linkage disequilibrium (LD) reference panel for the score construction step.^38^ Approximately 1.2 million common (MAF ≥1%) variants in HapMap3 were kept for score construction, as suggested in Vilhjálmsson et al.^39,40^ We computed PRS in three ways: LDpred, pruning and threshold (P+T), and a simple weighted combination of SNPs that reached genome-wide significance in our meta-analysis combining UK Biobank and CKDGen, a special case of P+T.

The primary PRS was calculated using the LDpred algorithm.^40^ For this method, we created 5 candidate LDpred PRSs under different assumptions for the fraction of causal variants. This Bayesian approach utilizes GWAS summary statistics to compute the posterior mean effect sizes for the genetic variants by assuming a prior of the joint effect sizes and incorporating the LD structure of the reference population. Two parameters of the LDpred need to be set by the users. One is the LD radius, which is the number of variants being adjusted for at each side of a variant. We set it to 400 (which corresponds to 1.2× 10^6^/3,000) based on Vilhjálmsson et al. The other parameter is the fraction of causal variants, ρ, which can be selected via parameter tuning on a separate dataset. Our tested ρ values were 1, 0.3, 0.1, 0.03, and 0.01, as suggested in Vilhjálmsson et al.^40^

We also implemented a second approach named pruning and thresholding. P+T scores were constructed with applying two filtering steps based on LD and *P* value.^41^ The variants are first pruned to only keep variants that have absolute pairwise correlation weaker than a threshold, *r*^2^ within certain genetic distance. The remaining variants are then filtered by removing the ones that have a *P*□value larger than a pre-defined threshold of significance. We created 30 candidate P+T PRSs based on four *r*^2^ levels (0.1, 0.2, 0.4, 0.6, and 0.8) and six *P*□values (5□×□10^−8^, 5□×□10^−6^, 5□×10^−4^, 0.05, 0.5, and 1). Finally, we created a “simple PRS” in a similar manner, using the most commonly used *r*^2^ level, 0.1, and *P*□value threshold, 5□×□10^−8^ (genome-wide significance).

For the PRS tuning, the 5 candidate LDpred PRSs, 30 candidate P+T PRSs, and one simple PRS were calculated in a tuning dataset of the remaining 10% unrelated EA participants in the UK Biobank (N=32,159). The best PRS of each approach was determined based on the proportion of the variance explained (R^2^) of eGFRcr that can be explained by the PRS. Specifically, we fitted a linear regression model with eGFRcr being the outcome, each candidate PRS being the exposure, and age at baseline and sex as the covariates. The best LDpred PRS and P+T PRS, as well as the simple PRS were carried forward into subsequent analyses in an independent validation dataset.

PRS validation was conducted in the 8,866 unrelated EA participants in ARIC. The R^2^ for eGFRcr by the best LDpred PRS, best P+T PRS, and simple PRS were calculated using the same approach with adjustment for the same covariates as in the tuning step. We compared the three PRSs with respect to number of SNPs included, phenotypic variance explained, and correlations with each other. In sensitivity analysis, we also directly implemented the PRS constructed and tuned on EA participants to the 2,871 unrelated AA participants in ARIC.

### Assessing Incident Kidney Diseases

Four incident kidney diseases were included in our study as outcomes: chronic kidney disease (CKD), end-stage kidney disease (ESKD), kidney failure, and acute kidney injury (AKI). CKD was defined based on the following criteria: eGFR <60 mL/min/1.73 m^2^ plus ≥30% eGFR decline during a follow-up visit comparing to baseline, ESKD cases identified through the direct linkage to the US Renal Data System (USRDS) registry, or International Classification of Diseases (ICD)-9/10-Clinical Modification (CM) codes **(Supplemental Table 1A)** representing CKD in any position of hospitalization or death records.^42^ ESKD was defined as having kidney transplant or dialysis in the USRDS registry. Kidney failure was defined by hospitalization codes (**Supplemental Table 1B**). AKI was defined by hospitalization or death codes (ICD-9-CM code: 584.X or ICD-10-CM code: N17.x).^43^

### Protein Measurements

Plasma proteins were measured in ARIC participants at visit 3 and visit 5 using the SOMAscan v.4 assay by SomaLogic. This platform uses Slow Off-rate Modified Aptamers (SOMAmers) to bind to targeted proteins and then uses DNA microarray to quantify them. SOMAscan v.4 includes 4,931 unique human proteins or protein complexes, with 95% of the proteins tagged by one modified aptamer and a total 5,211 modified aptamers. Protein measurements were reported as relative fluorescence units (RFUs).^12^ There were no missing values in the proteomic data.

Details of the quality control of the proteins were described elsewhere.^20^ Previous studies of SOMAscan v.3 consisting of 4,001 aptamers have shown high precision of this assay in quantifying proteins with median coefficient of variance (CV) of 4%-8%.^44-46^ In the current study, all proteins were log2 transformed.

### Assessing Covariates

Information on age, sex, center, and education level were assessed at baseline and current smoking status was assessed at all visits using an interviewer-administered questionnaires.^23^ Body mass index (BMI) was calculated as weight (in kilograms) divided by the square of height (in meters), both of which were measured at all visits. Clinical factors included history of hypertension, diabetes, and coronary heart disease (CHD). Hypertension was defined at all visits as systolic blood pressure ≥ 140 mm Hg, diastolic blood pressure ≥ 90 mm Hg, or use of antihypertensive medication in the past 2 weeks. Diabetes was defined at all visits as fasting blood glucose ≥ 126 mg/dL, non-fasting glucose ≥ 200 mg/dL, self-reported doctor-diagnosed diabetes, or use of diabetes medication in the past 2 weeks. CHD was defined at all visits as prior myocardial infarction (MI) observed on ECG, self-reported doctor-diagnosed heart attack, or self-reported cardiovascular surgery or coronary angioplasty. Albumin to creatinine ratio (ACR) was calculated as urinary albumin divided by urinary creatinine, with albumin being measured by an immunoturbidimetric method and creatinine being measured by a modified kinetic Jaffé method.

### Statistical Analysis

Baseline characteristics of the primary study population were examined. The R^2^ for eGFRcr at all visits except for visit 7 by the LDpred PRS, P+T PRS, and simple PRS were calculated as 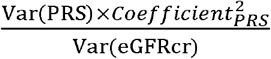 in a linear regression model of eGFRcr adjusting for age at the corresponded visit, sex, center, and first 10 genetic principal components (PCs). We also calculated the R^2^ at all visits with data for eGFRcys (visit 2 to visit 6) and ACR (visit 4 to visit 6). For comparison, we calculated the R^2^ for eGFRcr, eGFRcys, and ACR by the PRS using the same methods among the 2,871 AA participants as a sensitivity analysis.

We evaluated the association between PRS and incident kidney diseases. Using Cox proportional hazard models, we estimated hazard ratios (HR) and associated 95% confidence intervals (CI) of PRSs (per 1 SD lower PRS) for incident kidney diseases outcomes. We considered time at risk to start at visit 1 (1987–1989) and continue until the event of interest, death, loss to follow-up, or the end of follow-up (December 31, 2018). We evaluated three models: Model 1, which included age, sex, center, and first 10 genetic PCs; Model 2, which additionally included education, baseline BMI, baseline smoking status, baseline history of hypertension, diabetes, and CHD. In sensitivity analysis, we evaluated additional adjustments for eGFRcr and for both eGFRcr and ACR. As ACR was first measured at visit 4, the time to event for this sensitivity analysis started at visit 4 and the baseline covariates were also assessed at this time. We also examined the associations between PRS and all-cause mortality as well as comorbidities including incident hypertension, diabetes, CHD, and heart failure. Time to incident kidney diseases was assessed among quartiles of PRS using proportional hazard models and displayed using Kaplan-Meier survival curves.

To evaluate the association between PRS for kidney function and proteomic measurements, we conducted linear regression of LDpred PRS on 4,877 proteins measured at visit 3 and visit 5 adjusting for age, sex, center, and first 10 genetic PCs. These estimates reflect the difference in each log(2) transformed protein per normalized SD-unit higher in PRS for kidney function. Given that multiple statistical tests were performed, we utilized a Bonferroni adjusted P-value threshold of 0.05/4,877 ≈ 1.2 × 10^−5^ to indicate evidence for significant associations. We identified proteins significantly associated with LDpred PRS at both visit 3 and visit 5. We then examined their correlations and associations with eGFRcr and eGFRcys at each visit through Pearson correlation matrix and linear regression of PRS on eGFRcr or eGFRcys with adjustment for age, sex, center, and first 10 genetic PCs. Then we made scatter plots of correlations between those proteins and eGFR at each visit against their Pearson correlations with PRS. Analyses used R version 3.6.2 software (The R Foundation), two-tailed P-values, as well as statistical significance level of P < 0.05 except for the identification of proteomic signals, which was P < 1.02 × 10^−5^.

## RESULTS

### Characteristics of Study Cohort

Our primary study population included 8,886 participants (mean age 54.3 years; 53% female). Around 40% of them received college or above education. At baseline, 25% were smokers; mean BMI was 27.0 kg/m^2^; and the percentage of participants with prevalent hypertension, diabetes, and CHD were 26.7%, 8.6%, and 5.1% respectively. Over 30 years of follow up, the number of incident chronic kidney disease (CKD), end stage kidney disease (ESKD), kidney failure, and acute kidney injury (AKI) were 2,959, 137, 470, and 1,723, respectively (**Table 1**).

**Table 1.**
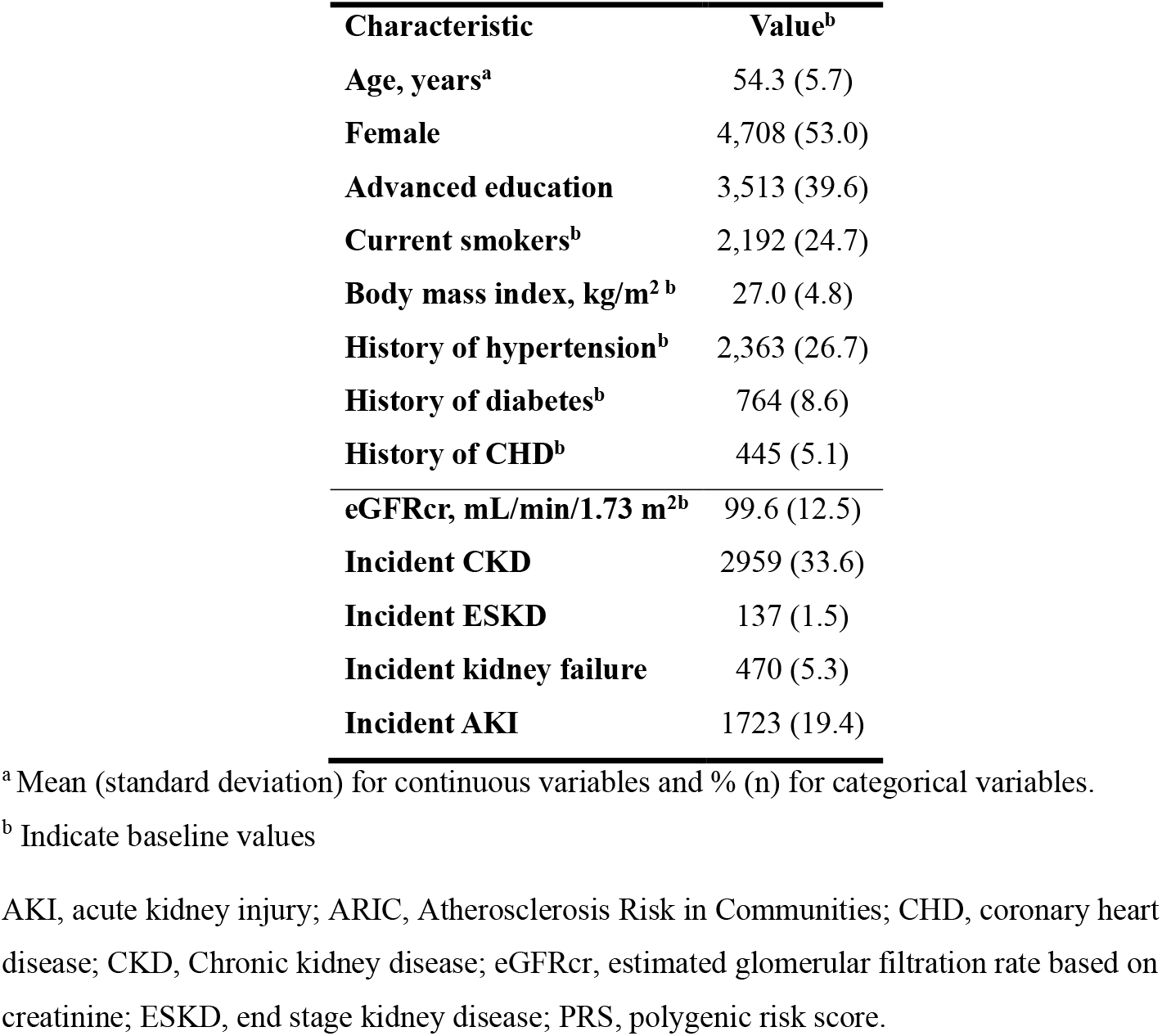
Characteristics of the study population in the Atherosclerosis Risk in Communities (ARIC) study (N=8,886).

### Characteristics of the Polygenic Risk Scores

LDpred PRS, P+T PRS, and simple PRS were all standardized to zero-mean and unit-variance and were approximately normally distributed in the population with. The technical details of the three PRSs are summarized in **Supplemental Table 2** and described in detail elsewhere.^47^ LDpred PRS was highly correlated with P+T PRS with a Pearson correlation coefficient (r) of 0.843, but moderately correlated with the simple PRS (r=0.580, **Supplemental Figure 2).** The adjusted eGFRcr variance explained by the LDpred PRS for kidney function was relatively consistent across the first four visits and slightly decreased at the last two visits, ranging from 5.4% to 8.7%. P+T PRS and simple PRS explained lower adjusted eGFRcr variance (P+T PRS: 4.5% to 7.2%; simple PRS: 3.4% to 5.6%). The adjusted eGFRcys variance was lower and its ranges for LDpred PRS, P+T PRS and simple PRS were 2.1% to 3.6%, 1.7% to 2.9%, and 1.2% to 2.3%, respectively. Variance explained for eGFR based on creatinine and cystatin (eGFRcr-cys) was intermediate and that for ACR was minimal. **(Supplemental Table 3).** As a comparison, directly applying the PRS trained and tuned on EA participants to AA participants led to substantially poorer score performance, with the eGFRcr variances explained by the LDpred PRS, P+T PRS, and simple PRS ranging from 1.4% to 2.2%, 0.5% to 1.4%, and 0.5% to 1.2% respectively (**Supplemental Table 4**).

### Associations between PRSs for Kidney Function and Incident Kidney Diseases

Categorizing the PRSs into deciles showed an incremental association with risk in Kaplan-Meier survival curves **(Figure 2).** In continuous analysis, we observed that the LDpred PRS for kidney function was strongly associated with all four incident kidney diseases: HRs (95% CI) per 1 SD-unit lower in LDpred PRS, indicating worse kidney function, were 1.33 (1.28, 1.39), 1.20 (1.00, 1.42), 1.17 (1.06, 1.28), and 1.07 (1.02, 1.12) for incident CKD, ESKD, kidney failure, and AKI respectively, after adjusting for age at baseline, sex, center, first 10 genetic PCs. Using P+T PRS and simple PRS, HRs for all incident kidney diseases were of smaller magnitude than using LDpred PRS and only statistically significant for CKD and kidney failure **(Table 2).**

**Table 2.**
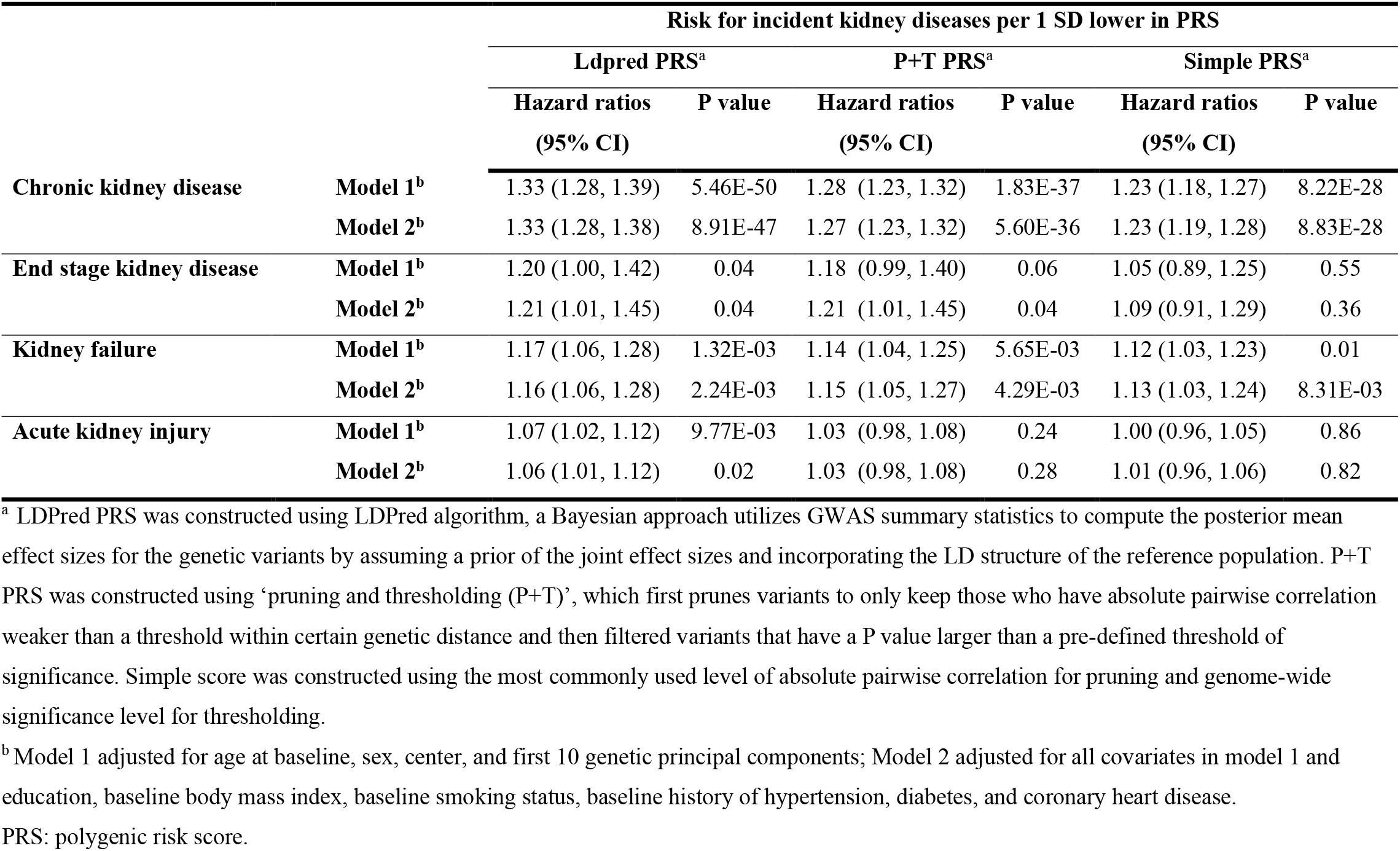
Risk for incident kidney diseases according to polygenic risk scores of kidney function (N=8,886).

|  |  | Risk for incident kidney diseases per 1 SD lower in PRS |  |  |  |  |  |
| --- | --- | --- | --- | --- | --- | --- | --- |
|  |  | Ldpred PRS<sup>a</sup> |  | P+T PRS<sup>a</sup> |  | Simple PRS<sup>a</sup> |  |
|  |  | Hazard ratios<br>(95% CI) | P value | Hazard ratios<br>(95% CI) | P value | Hazard ratios<br>(95% CI) | P value |
| Chronic kidney disease | Model 1<sup>b</sup> | 1.33 (1.28, 1.39) | 5.46E-50 | 1.28 (1.23, 1.32) | 1.83E-37 | 1.23 (1.18, 1.27) | 8.22E-28 |
|  | Model 2<sup>b</sup> | 1.33 (1.28, 1.38) | 8.91E-47 | 1.27 (1.23, 1.32) | 5.60E-36 | 1.23 (1.19, 1.28) | 8.83E-28 |
| End stage kidney disease | Model 1<sup>b</sup> | 1.20 (1.00, 1.42) | 0.04 | 1.18 (0.99, 1.40) | 0.06 | 1.05 (0.89, 1.25) | 0.55 |
|  | Model 2<sup>b</sup> | 1.21 (1.01, 1.45) | 0.04 | 1.21 (1.01, 1.45) | 0.04 | 1.09 (0.91, 1.29) | 0.36 |
| Kidney failure | Model 1<sup>b</sup> | 1.17 (1.06, 1.28) | 1.32E-03 | 1.14 (1.04, 1.25) | 5.65E-03 | 1.12 (1.03, 1.23) | 0.01 |
|  | Model 2<sup>b</sup> | 1.16 (1.06, 1.28) | 2.24E-03 | 1.15 (1.05, 1.27) | 4.29E-03 | 1.13 (1.03, 1.24) | 8.31E-03 |
| Acute kidney injury | Model 1<sup>b</sup> | 1.07 (1.02, 1.12) | 9.77E-03 | 1.03 (0.98, 1.08) | 0.24 | 1.00 (0.96, 1.05) | 0.86 |
|  | Model 2<sup>b</sup> | 1.06 (1.01, 1.12) | 0.02 | 1.03 (0.98, 1.08) | 0.28 | 1.01 (0.96, 1.06) | 0.82 |

<sup>a</sup> LDPred PRS was constructed using LDPred algorithm, a Bayesian approach utilizes GWAS summary statistics to compute the posterior mean effect sizes for the genetic variants by assuming a prior of the joint effect sizes and incorporating the LD structure of the reference population. P+T PRS was constructed using ‘pruning and thresholding (P+T)’, which first prunes variants to only keep those who have absolute pairwise correlation weaker than a threshold within certain genetic distance and then filtered variants that have a P value larger than a pre-defined threshold of significance. Simple score was constructed using the most commonly used level of absolute pairwise correlation for pruning and genome-wide significance level for thresholding.
<sup>b</sup> Model 1 adjusted for age at baseline, sex, center, and first 10 genetic principal components; Model 2 adjusted for all covariates in model 1 and education, baseline body mass index, baseline smoking status, baseline history of hypertension, diabetes, and coronary heart disease.
PRS: polygenic risk score.

**Figure 1.**
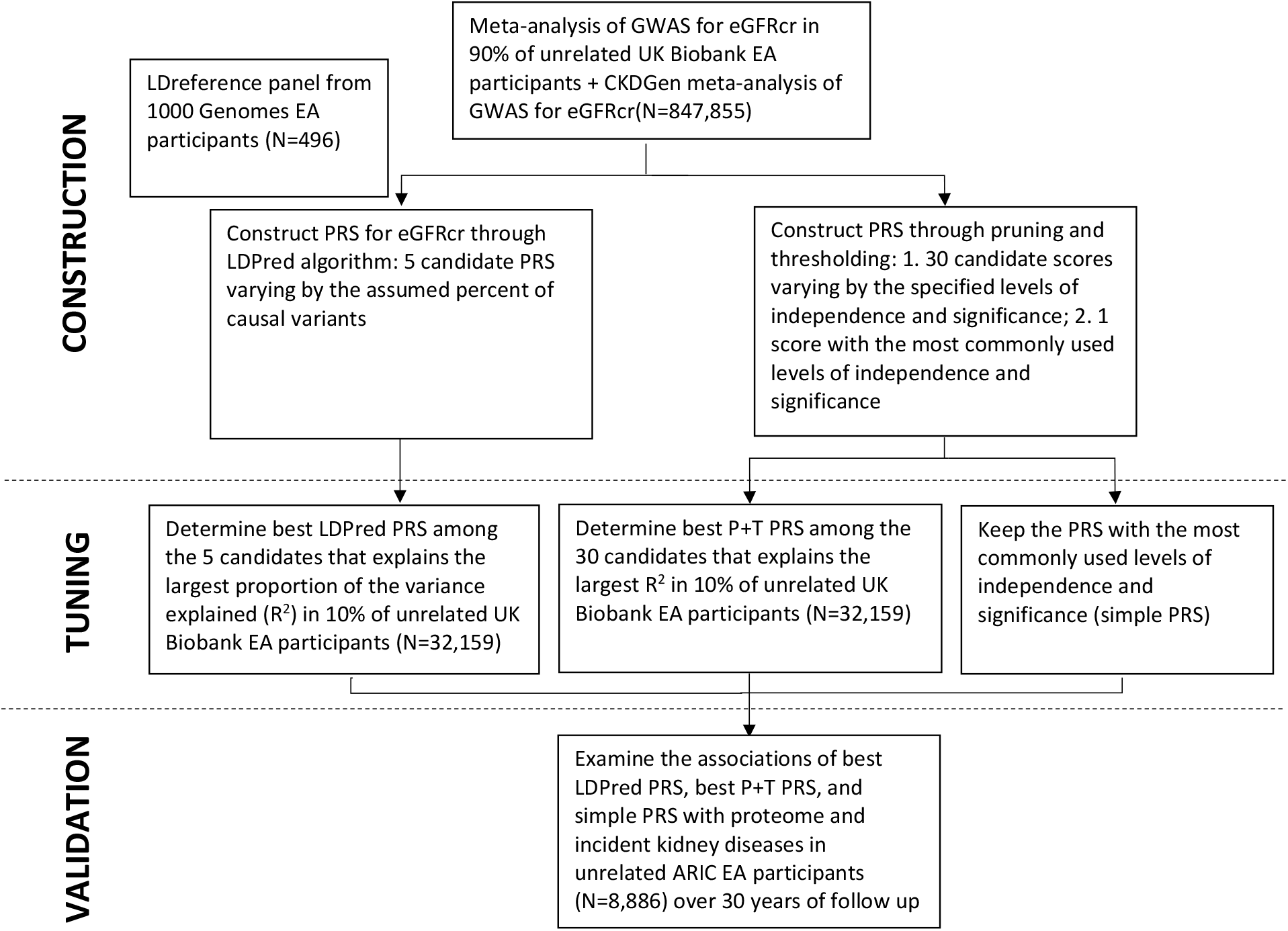
Study design and workflow. Three polygenic risk scores (PRS) for kidney function measured as estimated glomerular filtration rate based on creatinine level (eGFRcr) were constructed by a European-ancestry (EA) meta-analysis of UK Biobank GWAS for eGFRcr (90% of the cohort) and a meta-analysis of GWAS for eGFRcr conducted by the CKDGen Consortium using LDpred algorithm, pruning and threshold (P+T), and a simple weighted combination of SNPs that reached genome-wide significance, followed by parameters tuned using data from the remaining 10% of UK Biobank EA participants, then tested for their associations with proteome and incident kidney diseases in the Atherosclerosis Risk in Communities (ARIC) study.

**Figure 2.**
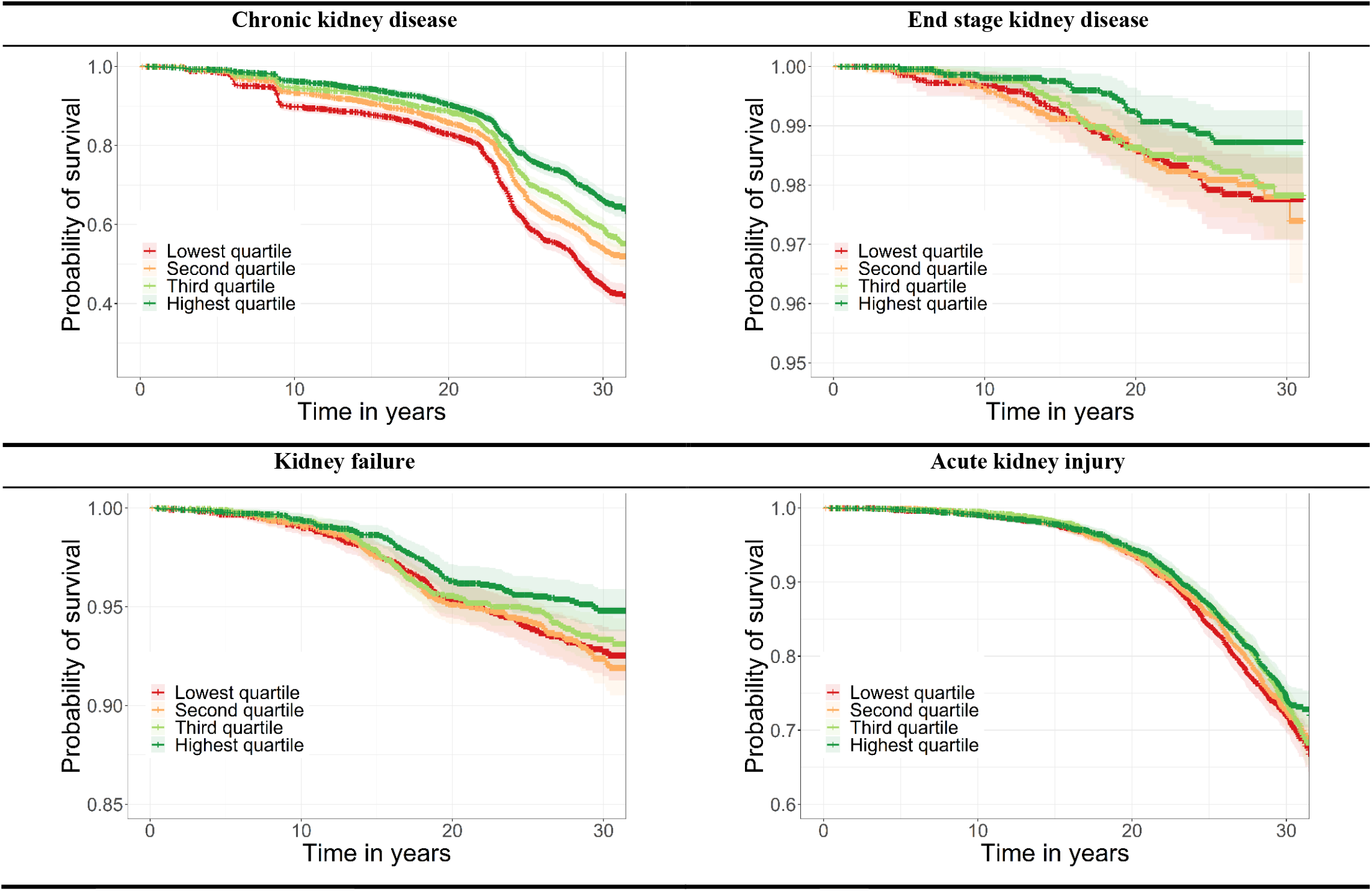
Association of quartiles of LDPred polygenic risk score of kidney function with incident kidney diseases (N=8,886). The LDPred polygenic risk score (PRS) for kidney function was categorized into quartiles, and was examined for their unadjusted associations with incident kidney diseases over 30-year of follow-up.

After adjustment for lifestyle and clinical risk factors (education, baseline BMI, baseline smoking status, and hypertension, diabetes, and CHD history at baseline) in Cox models, we observed limited changes in the risk estimates of the PRSs for all incident kidney diseases. However, risk estimates were substantially attenuated after additionally adjusting for the mediator eGFRcr **(Supplemental Table 5)**. Sensitivity analyses showed that additional adjustment for ACR made little difference **(Supplemental Table 6)** and the PRS did not associate with all-cause mortality as well as comorbidities (data not shown).

### Plasma Proteome as An Intermediate Trait

Using linear regression models adjusted for age, sex, center, and first 10 genetic PCs, we observed that 183 proteins were associated with LDpred PRS for kidney function at P = 1.2×10^−5^ level among 7,213 participants with valid proteomic measurements at visit 3, and 138 proteins among 3,666 participants at visit 5. Among those proteins, 108 were significant at both visits, which are 20 years apart. The strongest associations were with cystatin-C, collagen alpha-1(XV) chain, and desmocollin-2. Collagen alpha-1 (XV) chain exhibited strong and consistent associations with both eGFRcr and eGFRcys with a magnitude similar to that of cystatin with eGFRcr. For the 108 proteins consistently associated with LDpred PRS for kidney function, all but two of the associations were negative, indicating higher protein levels at lower kidney function. Testican 2 and angiostatin were the only two proteins with significant positive correlation to kidney function. The correlations with eGFRcr and eGFRcys measured at the corresponding visits were much stronger than those with the LDpred PRS, especially at visit 5 **(Figure 3, Supplemental Table 7,** median negative protein correlations at visit 5 of −0.0855, −0.4668, and −0.4697 with LDpred PRS, eGFRcr and eGFRcys with corresponding values at visit 3 of −0.0679, −0.2639 and −0.2820). This was also true for the significant positive correlations; the Pearson correlation coefficients with PRS, eGFRcr, and eGFRcys at visit3 were 0.100, 0.195, 0.197 for testican 2 and 0.067, 0.167, 0.258 for angiostatin respectively, and 0.103, 0.398, 0.433 for testican 2 and 0.095, 0.273, 0.344 for angiostatin respectively at visit 5.

**Figure 3.**
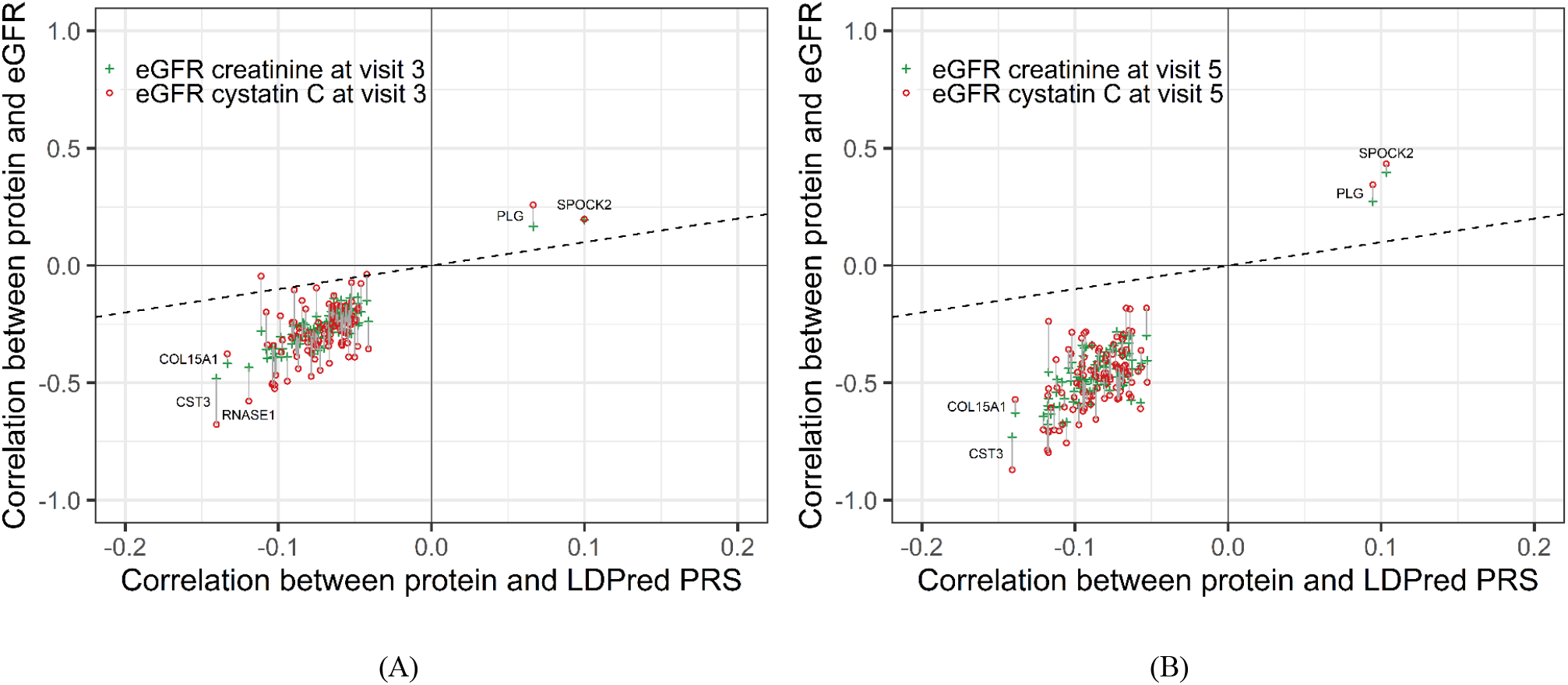
Scatter plots of Pearson’s correlations between protein and LDPred polygenic risk score for kidney function and correlation between protein and estimated glomerular filtration rate. Both protein measures and eGFR are visit specific (panel A – visit 3; panel B – visit 5). A total of 108 proteins were identified as significantly (Bonferroni threshold p < 1.02 × 10^−5^) associated with LDPred PRS at both visit 3 and visit 5 through linear regression of LDPred PRS on 4,877 proteins adjusting for age at the corresponded visits, sex, center, and first 10 genetic principal components. Visit 3 (N=7,213) was conducted during 1993-1995 when the mean age of study population was 60.4 years and visit 5 (N=3,666) was conducted during 2011-2013 when the mean age of study population was 75.9 years. The dashed line in grey is the identity line. COL15A1: collagen alpha-1(XV) chain; CST3: cystatin-C; PLG: angiostatin; RNASE1: ribonuclease pancreatic; SPOCK2: testican-2.

## DISCUSSION

In this community-based deeply phenotyped cohort of 8,866 middle-aged adults, we leveraged large studies to construct a range of polygenic risk scores for kidney function. A genome-wide score that included a weighted average of 1.2 million SNPs (LDpred PRS) showed the strongest association with kidney function while narrower risk scores (P+T PRS and simple PRS) showed weaker associations. The kidney function LDpred PRS was also associated with a range of plasma protein levels in mid-life and older age. However, these associations were weaker than the protein associations with eGFR itself. Genetic susceptibility to lower kidney function was associated with higher level of all significant proteins except testican 2 and angiostatin, which had lower levels at lower kidney function. Examination of future kidney disease progression showed genetic susceptibility to lower kidney function was associated with higher risk of CKD, ESKD, kidney failure, and AKI. LDpred PRS showed the strongest association, which was mediated by eGFR and independent of all other measured clinical and lifestyle risk factors including albuminuria.

An advantage of PRS for kidney function is that it can be assessed at any time, well before the emergence of lifestyle and clinical risk factors, such as elevated BMI, hypertension, and diabetes. Our results demonstrate that, for a spectrum of kidney diseases, not only diseases with established high heritability, but also entities like AKI whose genetic basis is less pronounced,^1-4,22^ PRS can now identify individuals with higher genetic risk for over 30 years of follow up, suggesting a potential role in further research and clinical medicine.

During the last decade, GWAS studies demonstrated thousands of genetic loci associated with hundreds of phenotypes.^48^ However, for most traits, the heritability explained by those SNPs (*h*^2^_gwas_) only explains a small portion of the estimated proportion of phenotypic variance due to additive genetic effects, i.e., narrow-sense heritability (*h*^2^).^49^ One of the proposed reasons for that was the existence of common causal variants of exceedingly low effect size which requires extremely large sample sizes to detect via GWAS.^50-54^ Using large studies for discovery and algorithms that incorporate variants across the genome, our results showed a significant improvement in the performance of PRS compared with previous efforts for score development (7% vs. 1.7 to 2.8% of variance explained).^6-11^ The phenotypic variance explained by the PRS was larger and in line with GWAS meta-analysis estimates.^5^ We also observed a statistically significant link between the genetic basis of eGFR and a spectrum of incident kidney disease outcomes.

In addition to the improved algorithm for constructing PRS, two other factors were important for improving the prediction performance of PRS. In recent years, mega cohorts and global genetics consortia provided sufficient power for detecting loci that confer only small changes in disease risk, which was a key factor in improving accuracy of PRS for disease prediction. Another factor is accurate identification of disease. In our study, we benefited from linkage to the USRDS registry and long term active follow-up for identifying cases of incident kidney diseases, which also improved the power of our analysis. As the incident kidney diseases were identified through different methods, the magnitude of the hazard ratios may also reflect outcome misclassification, which is lower for ESKD and higher for AKI.

Differences in plasma protein levels can provide clues for intermediate pathways between genetic susceptibility and disease. These alterations can happen as a result of the genetic susceptibility itself or secondary to other physiologic changes, including reduced eGFR. Our finding that many proteins were negatively associated with kidney function suggests that low renal clearance may result in relative protein accumulation. Increased protein levels in the setting of kidney disease may also reflect ongoing pathological processes, such as inflammation. Specific proteins are of interest. Collagen alpha-1 (XV) chain showed strong and consistent associations with PRS and both eGFRcr and eGFRcys with a magnitude similar to that of cystatin, the best marker for kidney function independent of demographics at current practice, suggesting its potential role as a marker for genetically predicted kidney function. It forms the alpha chain of type XV collagen, a structural component widely expressed across tissues, including kidney, with a predominant localization in the basement membrane zones.^55,56^ Experimental studies conducted on fetal kidneys demonstrated the existence of both its mRNAs and proteins at glomeruli and collecting ducts, as well as elevated levels in samples collected from patients with glomerular diseases.^57,58^ Testican 2, one of two proteins that was lower at lower kidney function, forms a structural component of the extracellular matrix through covalently binding with glycosaminoglycans.^59^ It is expressed in multiple tissues including the kidney and genetic variants in its gene, *SPOCK2*, have been strongly associated with bronchopulmonary dysplasia but the connection to kidney disease is largely unknown. This gene is not at or near any eGFR loci reported by previous GWAS results.^5^ Angiostatin, the other protein that was lower with lower kidney function, is a potent angiogenesis inhibitor generated through the proteolysis of plasminogen. Evidence also suggests its anti-inflammatory roles through hindering the recruitment of leukocyte^60^ and the movement of neutrophil and macrophage.^61,62^ Alterations in angiogenesis and inflammation have important roles in kidney disease pathophysiology.^63,64^ Previous animal experiments demonstrated that angiostatin overexpression slowed the progression of renal disease after chronic kidney injury and its decrease expression accelerated the pathogenesis process of diabetic nephropathy^65,66^ Observational studies have suggested elevated urinary angiostatin as potential biomarker of the disease severity and progression for IgA nephropathy and lupus nephritis.^67,68^

An interesting finding regarding the genetic and environmental influences on protein levels was that proteins’ correlations with kidney function were much stronger than those with PRS and that the difference was especially pronounced at visit 5 when the participants were in their 70s or 80s. It may suggest a decrease in genetic influence on human body with aging, which was also in line with our other observations that the phenotypic variance of kidney function explained by PRS were smaller at the last two visits comparing with earlier visits.

Our study had limitations. The PRS developed in our study were constructed, tuned, and validated in EA participants only, mirroring most genetic studies done so far. Since LD patterns, minor allele frequencies, effect sizes of common variants, and phenotypic features vary by ancestry, our PRS constructed based on GWAS results and LD structure of EA individuals will have poor disease prediction for individuals with other ancestries.^69,70^ When directly applying our PRS trained and tuned on EA participants to AA participants, the phenotypic variance explained was several-fold lower. It is therefore necessary and important for future efforts to include more multi-ethnic participants in genomic studies and develop novel methods that appropriately tailor genetic risk score to each ethnic group. The PRS presented in this study were for eGFRcr which means they may include genetic influences of creatinine metabolism as well as kidney function. However, we included eGFRcys as an outcome to assess the extent to which associations were robust to the kidney function marker used. Finally, we focused on common genetic variants in calculating the PRS recognizing future work may include additional variants, including rare variants with larger effects.

In conclusion, our results show polygenic risk scores for kidney function are associated with future risk of incident kidney diseases, including CKD progression, end-stage kidney disease, kidney failure, and acute kidney injury, over 30 years of follow up in a community-based cohort. This association was independent of most risk factors, including albuminuria, but was largely mediated through kidney function itself. A large number of plasma protein levels were elevated among individuals with high genetic risk for low kidney function while two proteins (testican-2 and angiostatin) had lower levels at lower kidney function. The protein associations were much stronger with concurrent kidney function than with PRS and were stronger at older age. Thus, progressive kidney disease may be a dominant influence on the proteome beyond eGFR genetic susceptibility.

## Acknowledgements

The authors thank the staff and participants of the ARIC study for their important contributions.

