## Supplemental Materials for "Polygenic Risk Scores for Kidney Function to the Circulating Proteome, and Incident Kidney Diseases: the Atherosclerosis Risk in Communities Study"

**Supplemental Figure 1. Flow chart of subjects included in the study.**

N=7,213

Study population for evaluating the association between PRS and proteomic measurements

N= 11,478

15,792 participants enrolled in the ARIC at baseline in 1987-1989

N= 8,886

Study population for evaluating the associations of PRS with estimated glomerular filtration rate (eGFR) and kidney diseases

Exclusion: 459 participants who were one of the close relative pairs (determined by the proportion of their genomes identical-by-state (IBS)).

Exclusion: 2,133 participants who did not have valid genetic information for creating polygenic risk scores (PRS).

N= 9,345

Exclusion: 4,314 participants whose self-reported race groups were not White

Exclusion: 1,673 participants without proteomic data at visit 3measurements.

**Supplemental Table 1. ICD-9/10 codes used for identifying chronic kidney disease and kidney failure.**

**(A) Chronic kidney disease (CKD)**

| **ICD-9-code** | **Description** | **ICD-10-code** |
| --- | --- | --- |
| 582 | Chronic glomerulonephritis | N03 |
| 583 | Nephritis and nephropathy |  |
| 585, 585.x where x≥3 | Chronic kidney disease | N18, N18.x where x≥3 |
| 586 | Renal failure | N19 |
| 587 | Renal sclerosis | N26 |
| 588 | Disorders resulting from impaired renal function | N25 |
| 403 | Hypertensive chronic kidney disease | I12 |
| 404 | Hypertensive heart and kidney disease | I13 |
| 593.9 | Unspecified disorder of the kidney and ureter |  |
| 250.4 | Diabetes with renal complications | E10.2, E11.2, E13.2 |
| V42.0 | Kidney replaced by transplant | Z94.0 |
| 55.6 | Transplant of kidney |  |
| 996.81 | Complications of transplanted kidney |  |
| V45.1^a^ | Renal dialysis status | Z99.2 |
| V56 ^a^ | Admission for dialysis treatment or session | Z49 |
| 39.95^a^ | Hemodialysis |  |
| 54.98^a^ | Peritoneal dialysis |  |
|  | Encounter for adjustment and management of vascular access device | Z45.2 |

^a^ Codes that are counted as incident CKD only if a concomitant acute kidney injury (AKI) code (ICD-9: 584.x, ICD-10: N17) is not present.

**(B) Kidney failure**

| **ICD-9-code** | **Description** | **ICD-10-code** |
| --- | --- | --- |
| V42.0 | Kidney replaced by transplant | Z94.0 |
| 55.6 | Transplant of kidney |  |
| 996.81 | Complications of transplanted kidney |  |
| V45.1^a^ | Renal dialysis status | Z99.2 |
| V56 ^a^ | Admission for dialysis treatment or session | Z49 |
| 39.95^a^ | Hemodialysis |  |
| 54.98^a^ | Peritoneal dialysis |  |
|  | Encounter for adjustment and management of vascular access device | Z45.2 |
| 585.5 | Chronic kidney disease stage 5 | N18.5 |
| 585.6 | End stage renal disease | N18.6 |
| 586 | Renal failure | N19 |
| 403.01 | Hypertensive chronic kidney disease, malignant, with CKD 5 or ESRD |  |
| 403.91 | Hypertensive chronic kidney disease, with CKD 5 or ESRD | I12.0 |

^a^ Codes that are not counted as incident kidney failure if: 1) for hospitalizations, a concurrent AKI code is present; 2) for deaths, if a concurrent AKI code is present without a concurrent CKD code.

**Supplemental Table 2. Technical details of polygenic risk scores.**

|  | **LDPred PRS** | **P+T PRS** | **Simple PRS** |
| --- | --- | --- | --- |
| **Features** | Entire summary results of variants with rescaling weights based on LD structure, effect size, and estimated causal fraction | r^2^ and P value thresholds to restrict variants | r^2^ and genome-wide significant threshold to restrict variants |
| **Settings** | LD structure from 1000 Genome  % of causal variants = 30% | $r^{2}$ = 0.1  P = 0.05 | $r^{2}$ = 0.1  P = 5 $*{10}^{-8}$ |
| **No. of candidate SNPs** | ~ 1.2 million | ~ 1.2 million | ~ 1.2 million |
| **No. of included SNPs** | ~ 1.2 million | 36,944 | 1,022 |
| **Rescaling weights** | Yes | No | No |

PRS: polygenic risk score.

| **P + T PRS**^a^ **vs. LDPred PRS**^a^ | **Simple PRS**^a^ **vs. LDPred PRS** | **Simple PRS vs. P + T PRS** |
| --- | --- | --- |
| 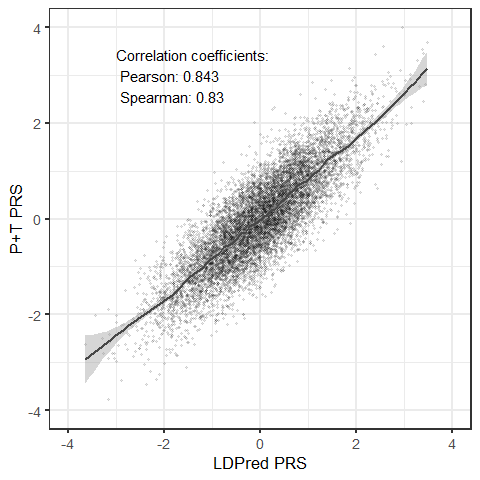 | 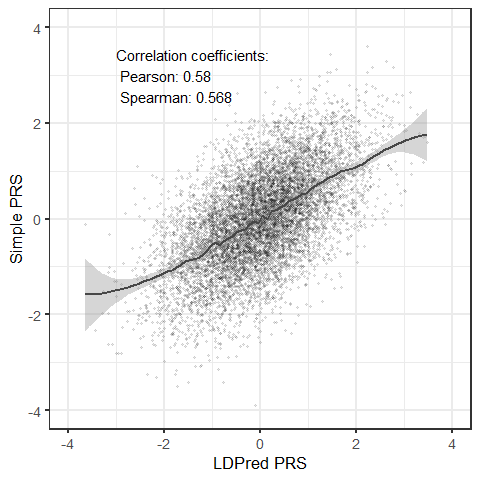 | 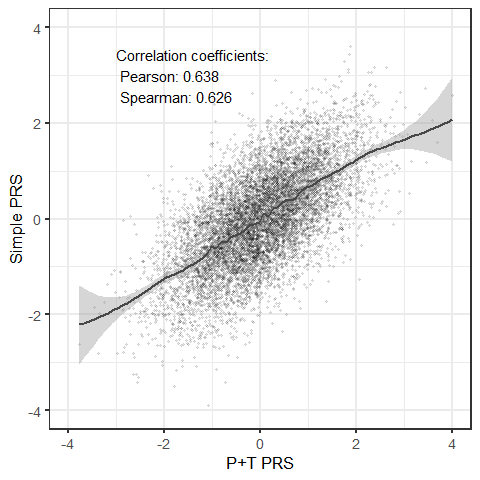 |

**Supplemental Figure 2.** **Scatter plots of polygenic risk scores (PRS) with locally weighted smoothing (LOESS) regression line.**

LDPred PRS was constructed using LDPred algorithm, a Bayesian approach utilizes GWAS summary statistics to compute the posterior mean effect sizes for the genetic variants by assuming a prior of the joint effect sizes and incorporating the LD structure of the reference population. P+T PRS was constructed using ‘pruning and thresholding (P+T)’, which first prunes variants to only keep those who have absolute pairwise correlation weaker than a threshold within certain genetic distance and then filtered variants that have a P value larger than a pre-defined threshold of significance. Simple score was constructed using the most commonly used level of absolute pairwise correlation for pruning and genome-wide significance level for thresholding.

**Supplemental Table 3. Adjusted proportion of the variance for estimated glomerular filtration rate (eGFR) and albumin to creatinine ratio (ACR) explained by polygenic risk scores (PRS) (N = 8,886).**

|  | **eGFRcr** | | | **eGFRcys** | | | **eGFRcr-cys** | | | **ACR** | | |
| --- | --- | --- | --- | --- | --- | --- | --- | --- | --- | --- | --- | --- |
|  | **LDPred PRS**^a^ | **P+T PRS**^a^ | **Simple PRS**^a^ | **LDPred PRS**^a^ | **P+T PRS**^a^ | **Simple PRS**^a^ | **LDPred PRS**^a^ | **P+T PRS**^a^ | **Simple PRS**^a^ | **LDPred PRS**^a^ | **P+T PRS**^a^ | **Simple PRS**^a^ |
| **Visit 1** | 0.070^b^ | 0.055 | 0.045 | - | - | - | - | - | - | - | - | - |
| **Visit 2** | 0.069 | 0.057 | 0.046 | 0.027 | 0.021 | 0.018 | 0.055 | 0.045 | 0.037 | - | - | - |
| **Visit 3** | 0.087 | 0.072 | 0.056 | 0.036 | 0.029 | 0.023 | 0.067 | 0.055 | 0.043 | - | - | - |
| **Visit 4** | 0.072 | 0.059 | 0.041 | 0.023 | 0.019 | 0.016 | 0.054 | 0.044 | 0.034 | 0.0005 | 0.002 | 0.001 |
| **Visit 5** | 0.054 | 0.045 | 0.034 | 0.021 | 0.017 | 0.012 | 0.038 | 0.031 | 0.022 | 0.002 | 0.002 | 0.001 |
| **Visit 6** | 0.058 | 0.050 | 0.041 | 0.025 | 0.023 | 0.017 | 0.042 | 0.038 | 0.030 | 0.0007 | 0.0002 | 0.0007 |

^a^ LDPred PRS was constructed using LDPred algorithm, a Bayesian approach utilizes GWAS summary statistics to compute the posterior mean effect sizes for the genetic variants by assuming a prior of the joint effect sizes and incorporating the LD structure of the reference population. P+T PRS was constructed using ‘pruning and thresholding (P+T)’, which first prunes variants to only keep those who have absolute pairwise correlation weaker than a threshold within certain genetic distance and then filtered variants that have a P value larger than a pre-defined threshold of significance. Simple score was constructed using the most commonly used level of absolute pairwise correlation for pruning and genome-wide significance level for thresholding.

^b^ Proportion of the variance for eGFR explained by PRS with adjusting for age at the corresponded visit, sex, center, and first 10 genetic principal components for all such values.

eGFRcr, estimated glomerular filtration rate based on creatinine; eGFRcr-cys, estimated glomerular filtration rate based on creatinine and cystatin; eGFRcysr, estimated glomerular filtration rate based on cystatin.

**Supplemental Table 4. Adjusted proportion of the variance for estimated glomerular filtration rate (eGFR) and albumin to creatinine ratio (ACR) explained by polygenic risk scores (PRS) among participants with African ancestry (N = 2,871).**

|  | **eGFRcr** | | | **eGFRcys** | | | **eGFRcr-cys** | | | **ACR** | | |
| --- | --- | --- | --- | --- | --- | --- | --- | --- | --- | --- | --- | --- |
|  | **LDPred PRS**^a^ | **P+T PRS**^a^ | **Simple PRS**^a^ | **LDPred PRS**^a^ | **P+T PRS**^a^ | **Simple PRS**^a^ | **LDPred PRS**^a^ | **P+T PRS**^a^ | **Simple PRS**^a^ | **LDPred PRS**^a^ | **P+T PRS**^a^ | **Simple PRS**^a^ |
| **Visit 1** | 0.010^b^ | 0.006 | 0.005 | - | - | - | - | - | - | - | - | - |
| **Visit 2** | 0.017 | 0.011 | 0.005 | 0.009 | 0.006 | 0.003 | 0.017 | 0.011 | 0.006 | - | - | - |
| **Visit 3** | 0.016 | 0.011 | 0.009 | 0.009 | 0.007 | 0.004 | 0.014 | 0.010 | 0.007 | - | - | - |
| **Visit 4** | 0.014 | 0.014 | 0.011 | 0.006 | 0.009 | 0.006 | 0.011 | 0.014 | 0.009 | 4.71E-06 | 0.0007 | 0.002 |
| **Visit 5** | 0.014 | 0.008 | 0.008 | 0.008 | 0.007 | 0.007 | 0.012 | 0.008 | 0.008 | 1.17 E-05 | 0.002 | 0.0006 |
| **Visit 6** | 0.022 | 0.005 | 0.012 | 0.008 | 0.003 | 0.010 | 0.015 | 0.004 | 0.012 | 0.0003 | 0.004 | 0.004 |

^a^ LDPred PRS was constructed using LDPred algorithm, a Bayesian approach utilizes GWAS summary statistics to compute the posterior mean effect sizes for the genetic variants by assuming a prior of the joint effect sizes and incorporating the LD structure of the reference population. P+T PRS was constructed using ‘pruning and thresholding (P+T)’, which first prunes variants to only keep those who have absolute pairwise correlation weaker than a threshold within certain genetic distance and then filtered variants that have a P value larger than a pre-defined threshold of significance. Simple score was constructed using the most commonly used level of absolute pairwise correlation for pruning and genome-wide significance level for thresholding.

^b^ Proportion of the variance for eGFR explained by PRS with adjusting for age at the corresponded visit, sex, center, and first 10 genetic principal components for all such values.

eGFRcr, estimated glomerular filtration rate based on creatinine; eGFRcr-cys, estimated glomerular filtration rate based on creatinine and cystatin; eGFRcysr, estimated glomerular filtration rate based on cystatin.

**Supplemental Table 5. Risk for incident kidney diseases adjusting for kidney fuction according to polygenic risk scores of kidney function (N=8,886).**

|  | **Risk for incident kidney diseases per 1 SD lower in PRS** | | | | | |
| --- | --- | --- | --- | --- | --- | --- |
|  | **Ldpred PRS**^a^ | | **P+T PRS**^a^ | | **Simple PRS**^a^ | |
|  | **Hazard ratios**  **(95% CI)** | **P value** | **Hazard ratios**  **(95% CI)** | **P value** | **Hazard ratios**  **(95% CI)** | **P value** |
| **Chronic kidney disease^b^** | 1.21 (1.16, 1.26) | 1.64E-19 | 1.17 (1.12, 1.21) | 3.42E-14 | 1.13 (1.09, 1.18) | 2.26E-10 |
| **End stage kidney disease^b^** | 0.98 (0.81, 1.18) | 0.81 | 1.01 (0.84, 1.22) | 0.90 | 0.91 (0.76, 1.08) | 0.29 |
| **Kidney failure^b^** | 1.03 (0.93, 1.14) | 0.57 | 1.00 (0.95, 1.05) | 0.91 | 1.03 (0.93, 1.13) | 0.59 |
| **Acute kidney injury^b^** | 1.03 (0.98, 1.08) | 0.30 | 1.04 (0.94, 1.14) | 0.49 | 0.98 (0.93, 1.03) | 0.35 |

^a^ LDPred PRS was constructed using LDPred algorithm, a Bayesian approach utilizes GWAS summary statistics to compute the posterior mean effect sizes for the genetic variants by assuming a prior of the joint effect sizes and incorporating the LD structure of the reference population. P+T PRS was constructed using ‘pruning and thresholding (P+T)’, which first prunes variants to only keep those who have absolute pairwise correlation weaker than a threshold within certain genetic distance and then filtered variants that have a P value larger than a pre-defined threshold of significance. Simple score was constructed using the most commonly used level of absolute pairwise correlation for pruning and genome-wide significance level for thresholding

^b^ Model adjusted for age at baseline, sex, center, first 10 genetic principal components, education, baseline body mass index, baseline smoking status, baseline history of hypertension, diabetes, and coronary heart disease, and estimated glomerular filtration rate based on creatinine (eGFRcr) measured at baseline.

PRS: polygenic risk score.

**Supplemental Table 6. Risk for incident kidney diseases according to polygenic risk scores of kidney function among participants who attended visit 4 (N=6,719).**

|  |  | **Risk for incident kidney diseases per 1 SD lower in PRS** | | | | | |
| --- | --- | --- | --- | --- | --- | --- | --- |
|  |  | **Ldpred PRS**^a^ | | **P+T PRS**^a^ | | **Simple PRS**^a^ | |
|  |  | **Hazard ratios**  **(95% CI)** | **P value** | **Hazard ratios**  **(95% CI)** | **P value** | **Hazard ratios**  **(95% CI)** | **P value** |
| **Chronic kidney disease** | **Model 1^b^** | 1.25 (1.20, 1.31) | 5.75E-22 | 1.21 (1.16, 1.27) | 5.21E-17 | 1.16 (1.11, 1.22) | 2.13E-11 |
|  | **Model 2^b^** | 1.25 (1.19, 1.31) | 1.90E-21 | 1.20 (1.15, 1.26) | 8.19E-16 | 1.17 (1.12, 1.22) | 3.64E-12 |
|  | **Model 3^b^** | 1.16 (1.10, 1.21) | 3.75E-09 | 1.12 (1.07, 1.17) | 2.91E-06 | 1.10 (1.05, 1.15) | 6.15E-05 |
|  | **Model 4^b^** | 1.16 (1.11, 1.22) | 1.25E-09 | 1.12 (1.07, 1.18) | 2.36E-06 | 1.11 (1.06, 1.16) | 1.55E-05 |
| **End stage kidney disease** | **Model 1^b^** | 1.17 (0.95, 1.43) | 1.32E-01 | 1.14 (0.93, 1.39) | 1.98E-01 | 0.99 (0.81, 1.2) | 8.83E-01 |
|  | **Model 2^b^** | 1.15 (0.94, 1.41) | 1.80E-01 | 1.14 (0.93, 1.40) | 2.10E-01 | 1.00 (0.82, 1.22) | 9.99E-01 |
|  | **Model 3^b^** | 0.78 (0.62, 0.97) | 2.84E-02 | 0.83 (0.66, 1.03) | 8.70E-02 | 0.79 (0.65, 0.97) | 2.06E-02 |
|  | **Model 4^b^** | 0.81 (0.65, 1.02) | 7.80E-02 | 0.85 (0.68, 1.06) | 1.43E-01 | 0.81 (0.66, 0.99) | 4.38E-02 |
| **Kidney failure** | **Model 1^b^** | 1.19 (1.07, 1.33) | 1.37E-03 | 1.17 (1.05, 1.31) | 3.51E-03 | 1.12 (1.00, 1.24) | 4.23E-02 |
|  | **Model 2^b^** | 1.19 (1.07, 1.33) | 1.98E-03 | 1.17 (1.05, 1.31) | 5.11E-03 | 1.11 (1.00, 1.24) | 4.69E-02 |
|  | **Model 3^b^** | 0.96 (0.85, 1.08) | 4.69E-01 | 0.97 (0.87, 1.09) | 6.16E-01 | 0.96 (0.86, 1.07) | 4.56E-01 |
|  | **Model 4^b^** | 0.98 (0.87, 1.10) | 7.06E-01 | 0.99 (0.88, 1.11) | 8.31E-01 | 0.98 (0.88, 1.09) | 7.16E-01 |
| **Acute kidney injury** | **Model 1^b^** | 1.07 (1.01, 1.13) | 1.51E-02 | 1.04 (0.99, 1.10) | 1.22E-01 | 1.01 (0.96, 1.07) | 6.13E-01 |
|  | **Model 2^b^** | 1.07 (1.02, 1.13) | 1.27E-02 | 1.04 (0.99, 1.10) | 1.58E-01 | 1.01 (0.96, 1.07) | 6.46E-01 |
|  | **Model 3^b^** | 1.00 (0.94, 1.06) | 8.92E-01 | 0.97 (0.92, 1.03) | 3.25E-01 | 0.96 (0.91, 1.01) | 1.02E-01 |
|  | **Model 4^b^** | 1.00 (0.94, 1.06) | 9.69E-01 | 0.97 (0.92, 1.03) | 3.35E-01 | 0.96 (0.91, 1.01) | 1.43E-01 |

^a^ LDPred PRS was constructed using LDPred algorithm, a Bayesian approach utilizes GWAS summary statistics to compute the posterior mean effect sizes for the genetic variants by assuming a prior of the joint effect sizes and incorporating the LD structure of the reference population. P+T PRS was constructed using ‘pruning and thresholding (P+T)’, which first prunes variants to only keep those who have absolute pairwise correlation weaker than a threshold within certain genetic distance and then filtered variants that have a P value larger than a pre-defined threshold of significance. Simple score was constructed using the most commonly used level of absolute pairwise correlation for pruning and genome-wide significance level for thresholding

^b^ Model 1 adjusted for age at visit 4, sex, center, and first 10 genetic principal components; Model 2 adjusted for all covariates in model 1 and education, body mass index at visit 4, smoking status at visit 4, history of hypertension, diabetes, and coronary heart disease at visit 4; Model 3 adjusted for all covariates in model 2 and albumin-to-creatinine ratio (ACR) measured at visit 4; Model 4 adjusted for all covariates in model 3 and estimated glomerular filtration rate based on creatinine (eGFRcr) measured at visit 4.

PRS: polygenic risk score

**Supplemental Table 7. Associations of LDPred polygenic risk score for kidney function and estimated glomerular filtration rate with proteins that are significantly associated with LDPred polygenic risk score at both visit 3 (N=7,213) and visit 5 (N = 3,666)^a^.**

**(A) LDPred PRS, eGFR measured at visit 3, and proteins measured at visit 3**

| **Gene** | **Full name** | **Visit 3 Protein and LDPred PRS^b,c^** | | | **Visit 3 Protein and Visit 3 eGFRcr ^c^** | | | **Visit 3 Protein and Visit 3 eGFRcys^c^** | | |
| --- | --- | --- | --- | --- | --- | --- | --- | --- | --- | --- |
|  |  | **Correlation**^e^ | **Beta (SE)** | **P** | **Correlation**^e^ | **Beta (SE)** | **P** | **Correlation**^e^ | **Beta (SE)** | **P** |
|  |  | **Positive significant correlations** | | | | | | | | |
| SPOCK2 | Testican-2 | 0.0999 | 0.0369 (0.0039) | 6.96E-21 | 0.1951 | 0.0054 (0.0003) | 8.77E-68 | 0.1972 | 0.0042 (0.0002) | 9.18E-71 |
| PLG | Angiostatin | 0.0665 | 0.0153 (0.0028) | 3.59E-08 | 0.1669 | 0.0021 (0.0002) | 8.06E-22 | 0.2579 | 0.0033 (0.0002) | 3.44E-86 |
| Positive significant correlations, median^f^ | | 0.0832 | | | 0.1810 | | | 0.2276 | | |
|  |  | **Negative significant correlations** | | | | | | | | |
| CST3 | Cystatin-C | -0.1405 | -0.0395 (0.003) | 7.62E-39 | -0.4811 | -0.009 (0.0002) | <2.23E-308 | -0.677 | -0.0104 (0.0001) | <2.23E-308 |
| COL15A1 | Collagen alpha-1(XV) chain | -0.1334 | -0.0351 (0.0029) | 4.28E-34 | -0.4171 | -0.0084 (0.0002) | <2.23E-308 | -0.3766 | -0.0058 (0.0002) | 2.50E-254 |
| RNASE1 | Ribonuclease pancreatic | -0.1193 | -0.0707 (0.0064) | 3.06E-28 | -0.4341 | -0.0166 (0.0005) | 2.00E-253 | -0.577 | -0.0179 (0.0003) | <2.23E-308 |
| DSC2 | Desmocollin-2 | -0.1031 | -0.0305 (0.003) | 2.07E-24 | -0.3896 | -0.0076 (0.0002) | 4.40E-240 | -0.3408 | -0.0049 (0.0002) | 1.20E-165 |
| COL6A3 | Collagen alpha-3(VI) chain | -0.104 | -0.0299 (0.0029) | 2.18E-24 | -0.3501 | -0.0064 (0.0002) | 1.30E-185 | -0.5068 | -0.0073 (0.0002) | <2.23E-308 |
| COL28A1 | Collagen alpha-1(XXVIII) chain | -0.1024 | -0.0321 (0.0032) | 8.27E-24 | -0.3534 | -0.0071 (0.0002) | 7.20E-181 | -0.5101 | -0.0081 (0.0002) | <2.23E-308 |
| CD59 | CD59 glycoprotein | -0.1072 | -0.031 (0.0031) | 1.09E-23 | -0.3941 | -0.0078 (0.0002) | 1.80E-234 | -0.3382 | -0.0049 (0.0002) | 1.70E-154 |
| TMED10 | Transmembrane emp24 domain-containing protein 10 | -0.1024 | -0.0359 (0.0036) | 2.63E-23 | -0.3768 | -0.0087 (0.0003) | 2.90E-213 | -0.5245 | -0.0095 (0.0002) | <2.23E-308 |
| GM2A | Ganglioside GM2 activator | -0.1016 | -0.0329 (0.0033) | 1.05E-22 | -0.3897 | -0.0082 (0.0002) | 3.40E-219 | -0.4669 | -0.0076 (0.0002) | <2.23E-308 |
| TNFRSF1A | Tumor necrosis factor receptor superfamily member 1A | -0.1038 | -0.0389 (0.004) | 2.43E-22 | -0.393 | -0.0095 (0.0003) | 4.10E-210 | -0.5008 | -0.0097 (0.0002) | <2.23E-308 |
| ART3 | Ecto-ADP-ribosyltransferase 3 | -0.1113 | -0.046 (0.0048) | 1.26E-21 | -0.2788 | -0.0101 (0.0004) | 2.30E-161 | -0.045 | -0.0015 (0.0003) | 4.25E-07 |
| CDNF | Cerebral dopamine neurotrophic factor | -0.1079 | -0.0286 (0.0031) | 1.54E-20 | -0.3566 | -0.0073 (0.0002) | 4.10E-209 | -0.1983 | -0.0029 (0.0002) | 6.99E-55 |
| SELM | Selenoprotein M | -0.098 | -0.0314 (0.0034) | 7.35E-20 | -0.3905 | -0.0081 (0.0003) | 2.40E-208 | -0.3691 | -0.0058 (0.0002) | 5.70E-174 |
| EFNB2 | Ephrin-B2 | -0.0973 | -0.0318 (0.0036) | 7.58E-19 | -0.3532 | -0.0082 (0.0003) | 1.70E-195 | -0.3181 | -0.0056 (0.0002) | 2.50E-151 |
| B2M | Beta-2-microglobulin | -0.0941 | -0.0274 (0.0032) | 8.66E-18 | -0.388 | -0.0074 (0.0002) | 1.50E-201 | -0.4931 | -0.0076 (0.0002) | <2.23E-308 |
| MYOC | Myocilin | -0.0986 | -0.0352 (0.0041) | 2.21E-17 | -0.3036 | -0.0082 (0.0003) | 1.40E-143 | -0.2149 | -0.0043 (0.0002) | 5.08E-67 |
| FABP4 | Fatty acid-binding protein, adipocyte | -0.0879 | -0.0522 (0.0062) | 5.45E-17 | -0.3044 | -0.0124 (0.0005) | 2.90E-145 | -0.3874 | -0.0121 (0.0004) | 7.20E-238 |
| LMAN2 | Vesicular integral-membrane protein VIP36 | -0.0884 | -0.0248 (0.003) | 9.20E-17 | -0.3218 | -0.0061 (0.0002) | 5.20E-155 | -0.3681 | -0.0054 (0.0002) | 1.40E-206 |
| MB | Myoglobin | -0.0895 | -0.0412 (0.005) | 1.35E-16 | -0.2509 | -0.0093 (0.0004) | 1.10E-127 | -0.1043 | -0.0029 (0.0003) | 4.92E-22 |
| IGFBP6 | Insulin-like growth factor-binding protein 6 | -0.0896 | -0.0231 (0.0028) | 2.04E-16 | -0.3139 | -0.0057 (0.0002) | 6.10E-150 | -0.2422 | -0.0032 (0.0002) | 8.14E-80 |
| LCN2 | Neutrophil gelatinase-associated lipocalin | -0.0901 | -0.0365 (0.0045) | 4.91E-16 | -0.2589 | -0.0074 (0.0003) | 5.99E-98 | -0.2976 | -0.0068 (0.0003) | 4.20E-139 |
| HSPB6 | Heat shock protein beta-6 | -0.0913 | -0.0384 (0.0047) | 6.33E-16 | -0.3341 | -0.0091 (0.0004) | 6.50E-134 | -0.3086 | -0.0063 (0.0003) | 5.10E-107 |
| MFAP2 | Microfibrillar-associated protein 2 | -0.0826 | -0.0262 (0.0033) | 2.19E-15 | -0.2509 | -0.0051 (0.0003) | 2.52E-85 | -0.2800 | -0.0043 (0.0002) | 6.80E-104 |
| FSTL3 | Follistatin-related protein 3 | -0.0726 | -0.0248 (0.0032) | 7.64E-15 | -0.3423 | -0.0065 (0.0002) | 8.10E-154 | -0.4465 | -0.0067 (0.0002) | 4.50E-286 |
| RGMB | RGM domain family member B | -0.0845 | -0.0212 (0.0027) | 8.31E-15 | -0.3045 | -0.0058 (0.0002) | 3.00E-165 | -0.1494 | -0.0021 (0.0002) | 2.02E-36 |
| ESAM | Endothelial cell-selective adhesion molecule | -0.091 | -0.0254 (0.0033) | 1.36E-14 | -0.2935 | -0.006 (0.0003) | 2.90E-119 | -0.2428 | -0.0039 (0.0002) | 6.59E-86 |
| CFD | Complement factor D | -0.0792 | -0.0256 (0.0033) | 1.45E-14 | -0.2749 | -0.0057 (0.0003) | 1.10E-107 | -0.2996 | -0.0046 (0.0002) | 2.00E-119 |
| JAM2 | Junctional adhesion molecule B | -0.0863 | -0.0202 (0.0026) | 2.38E-14 | -0.2633 | -0.0043 (0.0002) | 2.81E-94 | -0.2662 | -0.0034 (0.0002) | 2.20E-99 |
| NBL1 | Neuroblastoma suppressor of tumorigenicity 1 | -0.0781 | -0.0311 (0.0041) | 3.11E-14 | -0.3766 | -0.0095 (0.0003) | 2.90E-201 | -0.3767 | -0.0073 (0.0002) | 7.00E-199 |
| SMOC2 | SPARC-related modular calcium-binding protein 2 | -0.082 | -0.0272 (0.0036) | 4.14E-14 | -0.2827 | -0.0066 (0.0003) | 3.10E-121 | -0.1853 | -0.0031 (0.0002) | 1.82E-44 |
| TNFRSF1B | Tumor necrosis factor receptor superfamily member 1B | -0.0785 | -0.0286 (0.0038) | 4.52E-14 | -0.3007 | -0.0069 (0.0003) | 5.30E-120 | -0.4723 | -0.0088 (0.0002) | <2.23E-308 |
| GAS1 | Growth arrest-specific protein 1 | -0.086 | -0.0256 (0.0034) | 7.70E-14 | -0.3326 | -0.0069 (0.0003) | 2.00E-146 | -0.2843 | -0.0043 (0.0002) | 3.87E-94 |
| COL18A1 | Endostatin | -0.0729 | -0.0262 (0.0035) | 1.40E-13 | -0.2665 | -0.0061 (0.0003) | 3.70E-108 | -0.2652 | -0.0045 (0.0002) | 4.13E-98 |
| UNC5B | Netrin receptor UNC5B | -0.0771 | -0.0257 (0.0035) | 1.80E-13 | -0.3227 | -0.0066 (0.0003) | 1.60E-132 | -0.3293 | -0.0052 (0.0002) | 2.60E-134 |
| PPIC | Peptidyl-prolyl cis-trans isomerase C | -0.0771 | -0.0262 (0.0036) | 1.85E-13 | -0.2719 | -0.0063 (0.0003) | 7.40E-114 | -0.3251 | -0.0059 (0.0002) | 2.10E-168 |
| RNASE6 | Ribonuclease K6 | -0.0762 | -0.0367 (0.005) | 2.36E-13 | -0.2812 | -0.0085 (0.0004) | 1.40E-105 | -0.3993 | -0.0097 (0.0003) | 2.00E-237 |
| GABARAP | Gamma-aminobutyric acid receptor-associated protein | -0.0869 | -0.0195 (0.0027) | 2.87E-13 | -0.3085 | -0.005 (0.0002) | 7.90E-126 | -0.4391 | -0.0058 (0.0001) | 6.80E-308 |
| UNC5C | Netrin receptor UNC5C | -0.0739 | -0.0243 (0.0034) | 4.32E-13 | -0.2470 | -0.0049 (0.0003) | 3.57E-78 | -0.2430 | -0.0036 (0.0002) | 7.68E-71 |
| PTGDS | Prostaglandin-H2 D-isomerase | -0.0742 | -0.0323 (0.0045) | 6.61E-13 | -0.3607 | -0.0093 (0.0003) | 5.10E-157 | -0.3399 | -0.0065 (0.0003) | 9.80E-131 |
| DNAJB12 | DnaJ homolog subfamily B member 12 | -0.0807 | -0.0313 (0.0044) | 7.55E-13 | -0.3190 | -0.0086 (0.0003) | 3.10E-143 | -0.3283 | -0.0068 (0.0003) | 4.40E-152 |
| WFDC1 | WAP four-disulfide core domain protein 1 | -0.0777 | -0.0298 (0.0042) | 7.86E-13 | -0.3137 | -0.0077 (0.0003) | 1.70E-125 | -0.3575 | -0.007 (0.0002) | 8.00E-177 |
| CRABP2 | Cellular retinoic acid-binding protein 2 | -0.0809 | -0.0327 (0.0046) | 8.24E-13 | -0.2800 | -0.0083 (0.0003) | 1.80E-119 | -0.3613 | -0.0084 (0.0003) | 1.30E-211 |
| CCL14 | C-C motif chemokine 14 | -0.0735 | -0.0391 (0.0055) | 9.77E-13 | -0.2645 | -0.0092 (0.0004) | 5.30E-104 | -0.3223 | -0.0085 (0.0003) | 1.70E-149 |
| MIA | Melanoma-derived growth regulatory protein | -0.0751 | -0.026 (0.0036) | 9.93E-13 | -0.216 | -0.0053 (0.0003) | 2.00E-77 | -0.0955 | -0.0015 (0.0002) | 4.48E-12 |
| RARRES2 | Retinoic acid receptor responder protein 2 | -0.0671 | -0.0266 (0.0038) | 1.69E-12 | -0.2127 | -0.0054 (0.0003) | 2.32E-73 | -0.3435 | -0.0067 (0.0002) | 2.00E-200 |
| EFNA5 | Ephrin-A5 | -0.0778 | -0.0211 (0.003) | 2.39E-12 | -0.3063 | -0.0058 (0.0002) | 1.60E-135 | -0.2857 | -0.004 (0.0002) | 5.20E-109 |
| ATOX1 | Copper transport protein ATOX1 | -0.0839 | -0.0348 (0.005) | 5.50E-12 | -0.2437 | -0.0073 (0.0004) | 1.26E-75 | -0.3091 | -0.008 (0.0003) | 2.80E-155 |
| UNC5B | Netrin receptor UNC5B | -0.0757 | -0.0252 (0.0037) | 6.46E-12 | -0.289 | -0.0062 (0.0003) | 2.40E-104 | -0.3195 | -0.0053 (0.0002) | 2.20E-130 |
| EFNA4 | Ephrin-A4 | -0.0751 | -0.0196 (0.0029) | 1.06E-11 | -0.2856 | -0.0052 (0.0002) | 4.20E-119 | -0.3333 | -0.0046 (0.0002) | 2.20E-164 |
| TAGLN | Transgelin | -0.07 | -0.0279 (0.0041) | 1.50E-11 | -0.3513 | -0.0075 (0.0003) | 2.80E-118 | -0.2919 | -0.0041 (0.0002) | 1.86E-59 |
| TWSG1 | Twisted gastrulation protein homolog 1 | -0.0663 | -0.0154 (0.0023) | 1.51E-11 | -0.2736 | -0.0038 (0.0002) | 1.00E-101 | -0.2726 | -0.0029 (0.0001) | 5.80E-101 |
| CD46 | Membrane cofactor protein | -0.0688 | -0.0181 (0.0027) | 2.16E-11 | -0.2421 | -0.0043 (0.0002) | 4.39E-90 | -0.2577 | -0.0034 (0.0002) | 2.76E-97 |
| TNFRSF1B | Tumor necrosis factor receptor superfamily member 1B | -0.0666 | -0.0256 (0.0038) | 2.19E-11 | -0.2628 | -0.006 (0.0003) | 2.29E-90 | -0.417 | -0.0078 (0.0002) | 3.70E-267 |
| AIF1L | Allograft inflammatory factor 1-like | -0.0737 | -0.0213 (0.0032) | 3.97E-11 | -0.3092 | -0.0054 (0.0002) | 3.60E-102 | -0.3205 | -0.0044 (0.0002) | 6.70E-113 |
| CD55 | Complement decay-accelerating factor | -0.0669 | -0.0183 (0.0028) | 8.47E-11 | -0.2368 | -0.0043 (0.0002) | 2.38E-85 | -0.1639 | -0.0021 (0.0002) | 6.61E-35 |
| NPDC1 | Neural proliferation differentiation and control protein 1 | -0.0654 | -0.0233 (0.0036) | 9.86E-11 | -0.1954 | -0.0044 (0.0003) | 1.04E-53 | -0.2183 | -0.0037 (0.0002) | 1.74E-64 |
| VWC2 | Brorin | -0.0685 | -0.0242 (0.0038) | 1.13E-10 | -0.2693 | -0.0064 (0.0003) | 1.10E-105 | -0.2584 | -0.0046 (0.0002) | 6.11E-90 |
| CALCOCO2 | Calcium-binding and coiled-coil domain-containing protein 2 | -0.0637 | -0.0191 (0.003) | 2.12E-10 | -0.1397 | -0.0032 (0.0002) | 2.06E-40 | -0.1282 | -0.0022 (0.0002) | 1.99E-33 |
| EPHA2 | Ephrin type-A receptor 2 | -0.0673 | -0.0237 (0.0037) | 2.24E-10 | -0.2754 | -0.0059 (0.0003) | 7.89E-91 | -0.3183 | -0.0054 (0.0002) | 2.70E-128 |
| EPHB6 | Ephrin type-B receptor 6 | -0.0618 | -0.0168 (0.0027) | 3.81E-10 | -0.2274 | -0.0044 (0.0002) | 1.51E-98 | -0.168 | -0.0024 (0.0002) | 2.98E-49 |
| ROR2 | Tyrosine-protein kinase transmembrane receptor ROR2 | -0.0661 | -0.0239 (0.0038) | 3.89E-10 | -0.286 | -0.0066 (0.0003) | 5.70E-110 | -0.2946 | -0.0052 (0.0002) | 1.90E-113 |
| WFDC2 | WAP four-disulfide core domain protein 2 | -0.054 | -0.0305 (0.0049) | 5.69E-10 | -0.2893 | -0.0078 (0.0004) | 7.60E-91 | -0.3899 | -0.0087 (0.0003) | 2.10E-194 |
| FABP3 | Fatty acid-binding protein, heart | -0.0562 | -0.0381 (0.0063) | 1.89E-09 | -0.1851 | -0.0095 (0.0005) | 1.34E-81 | -0.2872 | -0.0106 (0.0004) | 4.00E-173 |
| DLK1 | Protein delta homolog 1 | -0.0645 | -0.0402 (0.0067) | 2.24E-09 | -0.2262 | -0.0093 (0.0005) | 5.42E-70 | -0.1857 | -0.0056 (0.0004) | 5.76E-43 |
| CD300C | CMRF35-like molecule 6 | -0.0591 | -0.0223 (0.0037) | 2.59E-09 | -0.148 | -0.0035 (0.0003) | 5.21E-32 | -0.2778 | -0.0053 (0.0002) | 4.50E-121 |
| EPHB4 | Ephrin type-B receptor 4 | -0.0635 | -0.0215 (0.0036) | 2.84E-09 | -0.2543 | -0.0056 (0.0003) | 4.14E-86 | -0.2586 | -0.0044 (0.0002) | 2.74E-89 |
| DLK1 | Protein delta homolog 1 | -0.0647 | -0.0374 (0.0063) | 3.05E-09 | -0.2185 | -0.0085 (0.0005) | 2.53E-66 | -0.1743 | -0.005 (0.0004) | 3.03E-38 |
| EFNA2 | Ephrin-A2 | -0.0662 | -0.0194 (0.0033) | 5.06E-09 | -0.2839 | -0.0058 (0.0003) | 5.90E-112 | -0.3221 | -0.0051 (0.0002) | 5.40E-145 |
| LRP10 | Low-density lipoprotein receptor-related protein 10 | -0.0699 | -0.016 (0.0027) | 5.89E-09 | -0.3042 | -0.005 (0.0002) | 1.70E-121 | -0.269 | -0.0033 (0.0002) | 3.97E-89 |
| IGFBP4 | Insulin-like growth factor-binding protein 4 | -0.0531 | -0.0204 (0.0035) | 1.02E-08 | -0.1375 | -0.0034 (0.0003) | 1.79E-34 | -0.2318 | -0.0044 (0.0002) | 1.62E-95 |
| SERPINF1 | Pigment epithelium-derived factor | -0.0612 | -0.0126 (0.0022) | 1.23E-08 | -0.207 | -0.0031 (0.0002) | 1.49E-73 | -0.2505 | -0.003 (0.0001) | 1.10E-113 |
| MXRA7 | Matrix-remodeling-associated protein 7 | -0.0606 | -0.0207 (0.0037) | 1.70E-08 | -0.2429 | -0.0056 (0.0003) | 1.92E-83 | -0.234 | -0.004 (0.0002) | 1.97E-73 |
| IL18BP | Interleukin-18-binding protein | -0.0585 | -0.0225 (0.004) | 1.82E-08 | -0.21 | -0.0044 (0.0003) | 1.99E-45 | -0.3404 | -0.0062 (0.0002) | 4.60E-152 |
| CAPG | Macrophage-capping protein | -0.0575 | -0.0335 (0.006) | 2.14E-08 | -0.1979 | -0.0065 (0.0005) | 4.18E-43 | -0.2807 | -0.0076 (0.0004) | 5.60E-100 |
| TNFRSF19 | Tumor necrosis factor receptor superfamily member 19 | -0.0604 | -0.0232 (0.0042) | 3.37E-08 | -0.2841 | -0.0074 (0.0003) | 6.90E-111 | -0.2617 | -0.0051 (0.0003) | 2.14E-89 |
| VIT | Vitrin | -0.0651 | -0.0222 (0.004) | 3.44E-08 | -0.2417 | -0.0057 (0.0003) | 7.07E-74 | -0.2456 | -0.0046 (0.0002) | 4.26E-81 |
| CPLX2 | Complexin-2 | -0.0561 | -0.0249 (0.0045) | 3.59E-08 | -0.2839 | -0.0081 (0.0003) | 8.50E-117 | -0.1717 | -0.0031 (0.0003) | 6.49E-29 |
| RETN | Resistin | -0.0623 | -0.0268 (0.0049) | 4.85E-08 | -0.2112 | -0.0068 (0.0004) | 1.81E-69 | -0.2422 | -0.006 (0.0003) | 8.50E-91 |
| ASGR1 | Asialoglycoprotein receptor 1 | -0.0567 | -0.0184 (0.0034) | 7.36E-08 | -0.2175 | -0.0048 (0.0003) | 1.34E-70 | -0.3015 | -0.0052 (0.0002) | 6.60E-142 |
| TNFRSF21 | Tumor necrosis factor receptor superfamily member 21 | -0.0538 | -0.0151 (0.0028) | 8.82E-08 | -0.1901 | -0.0036 (0.0002) | 2.12E-58 | -0.2484 | -0.0035 (0.0002) | 6.94E-96 |
| DCLK1 | Serine/threonine-protein kinase DCLK1 | -0.0634 | -0.022 (0.0041) | 9.64E-08 | -0.1991 | -0.0054 (0.0003) | 6.46E-63 | -0.2091 | -0.0044 (0.0002) | 3.35E-69 |
| PXDN | Peroxidasin homolog | -0.0586 | -0.0493 (0.0093) | 1.09E-07 | -0.2027 | -0.011 (0.0007) | 4.03E-51 | -0.3277 | -0.0149 (0.0005) | 1.50E-160 |
| EPHA1 | Ephrin type-A receptor 1 | -0.0521 | -0.0314 (0.0059) | 1.21E-07 | -0.1813 | -0.0066 (0.0005) | 1.39E-44 | -0.2125 | -0.006 (0.0004) | 5.37E-62 |
| PI3 | Elafin | -0.0493 | -0.0319 (0.0061) | 1.46E-07 | -0.2037 | -0.008 (0.0005) | 6.52E-63 | -0.2305 | -0.0071 (0.0004) | 2.58E-84 |
| SRL | Sarcalumenin | -0.0595 | -0.0252 (0.0048) | 1.65E-07 | -0.1838 | -0.0056 (0.0004) | 2.50E-49 | -0.1708 | -0.0039 (0.0003) | 1.96E-41 |
| IGFLR1 | IGF-like family receptor 1 | -0.05 | -0.0278 (0.0053) | 1.97E-07 | -0.252 | -0.0079 (0.0004) | 7.49E-80 | -0.3905 | -0.01 (0.0003) | 1.30E-222 |
| DLK2 | Protein delta homolog 2 | -0.0525 | -0.0227 (0.0044) | 2.12E-07 | -0.2905 | -0.0078 (0.0003) | 7.20E-117 | -0.2236 | -0.0044 (0.0003) | 2.07E-62 |
| VASN | Vasorin | -0.0643 | -0.0138 (0.0027) | 2.35E-07 | -0.1882 | -0.0033 (0.0002) | 7.55E-57 | -0.2238 | -0.0031 (0.0002) | 1.13E-85 |
| CLMP | CXADR-like membrane protein | -0.05 | -0.0228 (0.0045) | 4.19E-07 | -0.2243 | -0.0067 (0.0003) | 2.64E-82 | -0.2388 | -0.0053 (0.0003) | 7.24E-87 |
| NRXN3 | Neurexin-3-beta | -0.0521 | -0.0204 (0.0041) | 4.75E-07 | -0.1902 | -0.0044 (0.0003) | 1.22E-43 | -0.154 | -0.0024 (0.0002) | 1.16E-22 |
| XXYLT1 | Xyloside xylosyltransferase 1 | -0.0576 | -0.0172 (0.0034) | 4.86E-07 | -0.2024 | -0.0043 (0.0003) | 7.71E-59 | -0.207 | -0.0033 (0.0002) | 8.24E-59 |
| MAP2K2 | Dual specificity mitogen-activated protein kinase kinase 2 | -0.0528 | -0.0298 (0.006) | 5.61E-07 | -0.2286 | -0.008 (0.0005) | 3.14E-66 | -0.2525 | -0.0069 (0.0004) | 1.28E-81 |
| DDOST | Dolichyl-diphosphooligosaccharide--protein glycosyltransferase 48 kDa subunit | -0.0571 | -0.0163 (0.0033) | 6.69E-07 | -0.2293 | -0.0046 (0.0003) | 1.12E-70 | -0.2301 | -0.0036 (0.0002) | 1.33E-72 |
| B4GALT1 | Beta-1,4-galactosyltransferase 1 | -0.0562 | -0.0118 (0.0024) | 8.96E-07 | -0.2378 | -0.0036 (0.0002) | 1.01E-81 | -0.1896 | -0.002 (0.0001) | 7.78E-45 |
| TREM1 | Triggering receptor expressed on myeloid cells 1 | -0.0547 | -0.0218 (0.0045) | 9.58E-07 | -0.2204 | -0.0054 (0.0003) | 2.07E-52 | -0.3302 | -0.0068 (0.0003) | 3.10E-144 |
| MANSC1 | MANSC domain-containing protein 1 | -0.0482 | -0.0184 (0.0038) | 1.05E-06 | -0.1609 | -0.0039 (0.0003) | 3.03E-39 | -0.1886 | -0.0035 (0.0002) | 1.15E-52 |
| IL15RA | Interleukin-15 receptor subunit alpha | -0.0481 | -0.0213 (0.0044) | 1.06E-06 | -0.2545 | -0.0061 (0.0003) | 7.24E-72 | -0.3445 | -0.0069 (0.0003) | 5.00E-154 |
| CD93 | Complement component C1q receptor | -0.0483 | -0.0154 (0.0032) | 1.09E-06 | -0.1341 | -0.0033 (0.0002) | 1.77E-39 | -0.1857 | -0.0034 (0.0002) | 2.93E-69 |
| EFEMP1 | EGF-containing fibulin-like extracellular matrix protein 1 | -0.0411 | -0.0141 (0.0029) | 1.11E-06 | -0.2385 | -0.0034 (0.0002) | 3.41E-49 | -0.3549 | -0.0044 (0.0002) | 2.20E-142 |
| AMBP | Alpha-1-microglobulin | -0.0542 | -0.0154 (0.0032) | 1.20E-06 | -0.2113 | -0.004 (0.0002) | 4.13E-57 | -0.2833 | -0.0043 (0.0002) | 7.60E-114 |
| TMPO | Lamina-associated polypeptide 2, isoforms beta/gamma | -0.0588 | -0.0253 (0.0053) | 1.92E-06 | -0.2184 | -0.0066 (0.0004) | 1.38E-56 | -0.336 | -0.0088 (0.0003) | 1.00E-171 |
| NEGR1 | Neuronal growth regulator 1 | -0.0423 | -0.0113 (0.0024) | 3.64E-06 | -0.1503 | -0.0025 (0.0002) | 1.51E-38 | -0.0375 | -0.0002 (0.0001) | 0.253267 |
| PENK | Proenkephalin-A | -0.0555 | -0.0379 (0.0082) | 4.19E-06 | -0.1669 | -0.007 (0.0006) | 4.52E-27 | -0.2399 | -0.0086 (0.0005) | 2.08E-67 |
| SPINK7 | Serine protease inhibitor Kazal-type 7 | -0.0521 | -0.0222 (0.0049) | 5.39E-06 | -0.1937 | -0.0068 (0.0004) | 1.47E-70 | -0.0726 | -0.002 (0.0003) | 2.67E-11 |
| SMOC1 | SPARC-related modular calcium-binding protein 1 | -0.0477 | -0.0118 (0.0026) | 7.27E-06 | -0.2433 | -0.0035 (0.0002) | 7.45E-65 | -0.1798 | -0.0017 (0.0002) | 4.78E-26 |
| CST2 | Cystatin-SA | -0.0461 | -0.0337 (0.0076) | 9.71E-06 | -0.198 | -0.0096 (0.0006) | 9.57E-58 | -0.0767 | -0.0022 (0.0005) | 1.47E-06 |
| Negative significant correlations, median^f^ | | -0.0679 | | | -0.2639 | | | -0.2820 | | |

**(B) LDPred PRS, eGFR measured at visit 5, and proteins measured at visit 5**

| **Gene** | **Full name** | **Visit 5 Protein and LDPred PRS^b,d^** | | | **Visit 5 Protein and Visit 5 eGFRcr^d^** | | | **Visit 5 Protein and Visit 5 eGFRcys^d^** | | |
| --- | --- | --- | --- | --- | --- | --- | --- | --- | --- | --- |
|  |  | **Correlation** | **Beta (SE)** | **P** | **Correlation** | **Beta (SE)** | **P** | **Correlation** | **Beta (SE)** | **P** |
|  |  | **Positive significant correlations** | | | | | | | | |
| SPOCK2 | Testican-2 | 0.1033 | 0.0411 (0.0066) | 7.20E-10 | 0.3976 | 0.0096 (0.0004) | 6.00E-122 | 0.4333 | 0.009 (0.0003) | 9.85E-147 |
| PLG | Angiostatin | 0.0946 | 0.0192 (0.0037) | 1.66E-07 | 0.2734 | 0.0033 (0.0002) | 7.74E-48 | 0.3442 | 0.0037 (0.0002) | 7.82E-79 |
| Positive significant correlations, median^f^ | | 0.0989 | | | 0.3355 | | | 0.3887 | | |
|  |  | **Negative significant correlations** | | | | | | | | |
| COL15A1 | Collagen alpha-1(XV) chain | -0.139 | -0.0456 (0.0048) | 2.18E-21 | -0.6275 | -0.0112 (0.0002) | <2.23E-308 | -0.5708 | -0.0091 (0.0002) | 1.93E-313 |
| CST3 | Cystatin-C | -0.141 | -0.0535 (0.0056) | 4.59E-21 | -0.7304 | -0.015 (0.0003) | <2.23E-308 | -0.8703 | -0.0159 (0.0002) | <2.23E-308 |
| DSC2 | Desmocollin-2 | -0.1157 | -0.0493 (0.0058) | 3.72E-17 | -0.6337 | -0.0133 (0.0003) | <2.23E-308 | -0.6052 | -0.0112 (0.0003) | 8.89E-323 |
| CD59 | CD59 glycoprotein | -0.1177 | -0.0433 (0.0051) | 5.41E-17 | -0.5965 | -0.0111 (0.0003) | 1.64E-304 | -0.5533 | -0.0091 (0.0002) | 1.11E-266 |
| TNFRSF1A | Tumor necrosis factor receptor superfamily member 1A | -0.1173 | -0.0563 (0.0067) | 9.16E-17 | -0.6328 | -0.0151 (0.0003) | <2.23E-308 | -0.71 | -0.0151 (0.0003) | <2.23E-308 |
| GM2A | Ganglioside GM2 activator | -0.1207 | -0.0542 (0.0065) | 1.34E-16 | -0.6421 | -0.015 (0.0003) | <2.23E-308 | -0.6991 | -0.0145 (0.0003) | <2.23E-308 |
| RNASE1 | Ribonuclease pancreatic | -0.118 | -0.104 (0.0126) | 2.51E-16 | -0.6764 | -0.0303 (0.0006) | <2.23E-308 | -0.7862 | -0.0313 (0.0004) | <2.23E-308 |
| COL6A3 | Collagen alpha-3(VI) chain | -0.1173 | -0.0483 (0.0059) | 3.33E-16 | -0.6145 | -0.0129 (0.0003) | 5.23E-317 | -0.7037 | -0.0132 (0.0002) | <2.23E-308 |
| TMED10 | Transmembrane emp24 domain-containing protein 10 | -0.1173 | -0.0596 (0.0073) | 4.59E-16 | -0.6933 | -0.018 (0.0003) | <2.23E-308 | -0.7963 | -0.0183 (0.0003) | <2.23E-308 |
| ART3 | Ecto-ADP-ribosyltransferase 3 | -0.1173 | -0.0564 (0.007) | 7.09E-16 | -0.4533 | -0.0128 (0.0004) | 4.54E-207 | -0.2375 | -0.0062 (0.0004) | 3.15E-60 |
| EFNB2 | Ephrin-B2 | -0.1172 | -0.0469 (0.0058) | 8.70E-16 | -0.5655 | -0.0118 (0.0003) | 4.93E-262 | -0.5251 | -0.0097 (0.0003) | 7.08E-230 |
| COL28A1 | Collagen alpha-1(XXVIII) chain | -0.1136 | -0.0492 (0.0062) | 2.47E-15 | -0.6049 | -0.0133 (0.0003) | 3.74E-300 | -0.7006 | -0.0138 (0.0002) | <2.23E-308 |
| NBL1 | Neuroblastoma suppressor of tumorigenicity 1 | -0.1082 | -0.0692 (0.0091) | 3.25E-14 | -0.6745 | -0.0218 (0.0004) | <2.23E-308 | -0.6762 | -0.019 (0.0004) | <2.23E-308 |
| MFAP2 | Microfibrillar-associated protein 2 | -0.1118 | -0.0406 (0.0054) | 6.35E-14 | -0.4859 | -0.009 (0.0003) | 2.45E-169 | -0.5207 | -0.0086 (0.0003) | 1.09E-206 |
| PXDN | Peroxidasin homolog | -0.1102 | -0.0894 (0.0119) | 8.25E-14 | -0.6003 | -0.0255 (0.0006) | 9.65E-299 | -0.7044 | -0.0265 (0.0005) | <2.23E-308 |
| CD55 | Complement decay-accelerating factor | -0.1042 | -0.0329 (0.0044) | 1.03E-13 | -0.438 | -0.0068 (0.0003) | 5.54E-143 | -0.3575 | -0.0049 (0.0002) | 3.66E-96 |
| TWSG1 | Twisted gastrulation protein homolog 1 | -0.1069 | -0.0295 (0.004) | 2.32E-13 | -0.5667 | -0.008 (0.0002) | 8.36E-250 | -0.6029 | -0.0074 (0.0002) | 4.31E-298 |
| FSTL3 | Follistatin-related protein 3 | -0.0975 | -0.0404 (0.0055) | 2.43E-13 | -0.5862 | -0.0112 (0.0003) | 1.06E-263 | -0.6781 | -0.0115 (0.0002) | <2.23E-308 |
| LMAN2 | Vesicular integral-membrane protein VIP36 | -0.1013 | -0.0407 (0.0056) | 2.86E-13 | -0.5818 | -0.0116 (0.0003) | 2.60E-282 | -0.6133 | -0.0109 (0.0002) | <2.23E-308 |
| ROR2 | Tyrosine-protein kinase transmembrane receptor ROR2 | -0.1006 | -0.0481 (0.0066) | 4.42E-13 | -0.5376 | -0.0123 (0.0004) | 3.76E-213 | -0.5614 | -0.0113 (0.0003) | 6.21E-247 |
| B2M | Beta-2-microglobulin | -0.1056 | -0.0452 (0.0062) | 4.69E-13 | -0.6675 | -0.0147 (0.0003) | <2.23E-308 | -0.7559 | -0.0147 (0.0002) | <2.23E-308 |
| MB | Myoglobin | -0.102 | -0.0543 (0.0075) | 6.77E-13 | -0.413 | -0.0126 (0.0004) | 1.27E-169 | -0.2849 | -0.008 (0.0004) | 2.07E-86 |
| LCN2 | Neutrophil gelatinase-associated lipocalin | -0.1087 | -0.0556 (0.0077) | 8.19E-13 | -0.4967 | -0.0139 (0.0004) | 1.12E-196 | -0.5418 | -0.0135 (0.0004) | 2.24E-258 |
| RGMB | RGM domain family member B | -0.1026 | -0.0306 (0.0043) | 8.48E-13 | -0.492 | -0.0075 (0.0002) | 2.41E-192 | -0.3779 | -0.0051 (0.0002) | 4.14E-110 |
| CDNF | Cerebral dopamine neurotrophic factor | -0.1123 | -0.0385 (0.0054) | 1.10E-12 | -0.5404 | -0.0107 (0.0003) | 4.47E-248 | -0.4012 | -0.0069 (0.0003) | 3.14E-127 |
| SELM | Selenoprotein M | -0.0965 | -0.0423 (0.0059) | 1.13E-12 | -0.6084 | -0.0126 (0.0003) | 1.41E-295 | -0.5979 | -0.0109 (0.0003) | 3.42E-292 |
| SRL | Sarcalumenin | -0.1026 | -0.0455 (0.0065) | 3.01E-12 | -0.458 | -0.0108 (0.0004) | 6.11E-167 | -0.3738 | -0.0078 (0.0003) | 3.17E-111 |
| IGFBP6 | Insulin-like growth factor-binding protein 6 | -0.0848 | -0.0269 (0.0038) | 3.15E-12 | -0.4328 | -0.0062 (0.0002) | 1.11E-154 | -0.3647 | -0.0047 (0.0002) | 8.52E-118 |
| EPHA2 | Ephrin type-A receptor 2 | -0.0931 | -0.043 (0.0064) | 2.64E-11 | -0.5039 | -0.011 (0.0004) | 4.60E-179 | -0.539 | -0.0105 (0.0003) | 7.15E-222 |
| TNFRSF1B | Tumor necrosis factor receptor superfamily member 1B | -0.0937 | -0.0481 (0.0072) | 3.21E-11 | -0.4956 | -0.0125 (0.0004) | 4.63E-183 | -0.609 | -0.0138 (0.0003) | 2.77E-322 |
| EFNA5 | Ephrin-A5 | -0.0966 | -0.0348 (0.0052) | 3.30E-11 | -0.5222 | -0.0096 (0.0003) | 7.39E-207 | -0.5068 | -0.0082 (0.0003) | 9.21E-199 |
| CD46 | Membrane cofactor protein | -0.0989 | -0.0289 (0.0043) | 3.32E-11 | -0.4917 | -0.0074 (0.0002) | 9.14E-176 | -0.5017 | -0.0066 (0.0002) | 3.77E-186 |
| COL18A1 | Endostatin | -0.0921 | -0.033 (0.0051) | 1.07E-10 | -0.5416 | -0.0096 (0.0003) | 3.16E-220 | -0.5732 | -0.0089 (0.0002) | 1.16E-256 |
| EFNA4 | Ephrin-A4 | -0.0904 | -0.033 (0.0051) | 1.16E-10 | -0.5355 | -0.0097 (0.0003) | 8.30E-225 | -0.5568 | -0.009 (0.0002) | 2.98E-262 |
| TNFRSF1B | Tumor necrosis factor receptor superfamily member 1B | -0.0864 | -0.0464 (0.0072) | 1.71E-10 | -0.5315 | -0.0135 (0.0004) | 2.83E-216 | -0.6549 | -0.0149 (0.0003) | <2.23E-308 |
| WFDC1 | WAP four-disulfide core domain protein 1 | -0.0947 | -0.0421 (0.0066) | 1.81E-10 | -0.5379 | -0.0123 (0.0004) | 1.08E-216 | -0.5405 | -0.0108 (0.0003) | 3.02E-224 |
| UNC5B | Netrin receptor UNC5B | -0.0855 | -0.0352 (0.0055) | 2.30E-10 | -0.5099 | -0.0097 (0.0003) | 1.73E-187 | -0.5222 | -0.0087 (0.0003) | 1.15E-205 |
| RETN | Resistin | -0.094 | -0.0449 (0.0071) | 2.36E-10 | -0.3516 | -0.0084 (0.0004) | 2.02E-82 | -0.3901 | -0.0085 (0.0004) | 1.30E-112 |
| TNFRSF21 | Tumor necrosis factor receptor superfamily member 21 | -0.0856 | -0.028 (0.0044) | 2.57E-10 | -0.4081 | -0.0063 (0.0003) | 3.85E-120 | -0.419 | -0.0057 (0.0002) | 5.35E-134 |
| MYOC | Myocilin | -0.0957 | -0.0406 (0.0064) | 3.24E-10 | -0.4303 | -0.0103 (0.0004) | 4.10E-155 | -0.3091 | -0.0066 (0.0003) | 1.50E-80 |
| CFD | Complement factor D | -0.08 | -0.0293 (0.0046) | 3.47E-10 | -0.4101 | -0.0066 (0.0003) | 2.02E-118 | -0.4065 | -0.0059 (0.0002) | 1.72E-124 |
| PTGDS | Prostaglandin-H2 D-isomerase | -0.0898 | -0.0407 (0.0065) | 4.88E-10 | -0.5897 | -0.0134 (0.0003) | 3.74E-273 | -0.5922 | -0.0119 (0.0003) | 1.74E-285 |
| CD300C | CMRF35-like molecule 6 | -0.0945 | -0.0392 (0.0063) | 5.02E-10 | -0.3838 | -0.0083 (0.0004) | 7.48E-103 | -0.4669 | -0.009 (0.0003) | 6.25E-167 |
| FABP4 | Fatty acid-binding protein, adipocyte | -0.0807 | -0.0524 (0.0084) | 6.21E-10 | -0.4937 | -0.0156 (0.0005) | 9.64E-212 | -0.5664 | -0.0156 (0.0004) | 5.28E-296 |
| TNFRSF19 | Tumor necrosis factor receptor superfamily member 19 | -0.0898 | -0.0458 (0.0074) | 7.25E-10 | -0.4861 | -0.0126 (0.0004) | 4.04E-175 | -0.4458 | -0.0101 (0.0004) | 3.27E-147 |
| ESAM | Endothelial cell-selective adhesion molecule | -0.0956 | -0.0296 (0.0048) | 8.90E-10 | -0.4823 | -0.0082 (0.0003) | 7.00E-178 | -0.4412 | -0.0065 (0.0002) | 1.19E-147 |
| UNC5C | Netrin receptor UNC5C | -0.0802 | -0.0343 (0.0056) | 1.04E-09 | -0.3872 | -0.0073 (0.0003) | 2.58E-100 | -0.4023 | -0.0068 (0.0003) | 1.00E-114 |
| SEMG2 | Protein delta homolog 1 | -0.0939 | -0.0594 (0.0097) | 1.04E-09 | -0.3515 | -0.0122 (0.0006) | 7.05E-92 | -0.2881 | -0.0087 (0.0005) | 2.54E-61 |
| DLK1 | Protein delta homolog 1 | -0.0929 | -0.0579 (0.0096) | 1.91E-09 | -0.3443 | -0.0118 (0.0006) | 7.20E-88 | -0.2819 | -0.0084 (0.0005) | 1.20E-58 |
| TREM1 | Triggering receptor expressed on myeloid cells 1 | -0.0869 | -0.0421 (0.007) | 2.09E-09 | -0.4217 | -0.0097 (0.0004) | 4.48E-113 | -0.468 | -0.0098 (0.0003) | 1.03E-155 |
| DCLK1 | Serine/threonine-protein kinase DCLK1 | -0.0877 | -0.0405 (0.0068) | 2.35E-09 | -0.4399 | -0.0106 (0.0004) | 1.78E-148 | -0.4479 | -0.0096 (0.0003) | 6.01E-163 |
| GAS1 | Growth arrest-specific protein 1 | -0.0863 | -0.0317 (0.0053) | 2.63E-09 | -0.4827 | -0.0087 (0.0003) | 1.22E-160 | -0.4712 | -0.0074 (0.0003) | 1.02E-153 |
| JAM2 | Junctional adhesion molecule B | -0.0984 | -0.0236 (0.004) | 2.91E-09 | -0.4757 | -0.0068 (0.0002) | 4.07E-180 | -0.455 | -0.0056 (0.0002) | 1.03E-160 |
| EPHB6 | Ephrin type-B receptor 6 | -0.0811 | -0.0272 (0.0046) | 2.95E-09 | -0.5061 | -0.0079 (0.0003) | 4.87E-185 | -0.4816 | -0.0065 (0.0002) | 2.32E-162 |
| MXRA7 | Matrix-remodeling-associated protein 7 | -0.0794 | -0.0377 (0.0065) | 7.36E-09 | -0.5043 | -0.0114 (0.0004) | 1.20E-188 | -0.4854 | -0.0096 (0.0003) | 1.52E-177 |
| PPIC | Peptidyl-prolyl cis-trans isomerase C | -0.0794 | -0.0344 (0.0059) | 8.20E-09 | -0.4335 | -0.0093 (0.0003) | 1.57E-147 | -0.4598 | -0.0089 (0.0003) | 1.81E-182 |
| MANSC1 | MANSC domain-containing protein 1 | -0.0796 | -0.0312 (0.0054) | 8.40E-09 | -0.3686 | -0.0067 (0.0003) | 3.27E-89 | -0.3798 | -0.0062 (0.0003) | 7.80E-103 |
| CCL14 | C-C motif chemokine 14 | -0.0881 | -0.0531 (0.0092) | 8.74E-09 | -0.4783 | -0.0153 (0.0005) | 7.84E-168 | -0.555 | -0.0159 (0.0004) | 3.89E-251 |
| DNAJB12 | DnaJ homolog subfamily B member 12 | -0.0882 | -0.0389 (0.0068) | 1.07E-08 | -0.5364 | -0.0128 (0.0004) | 5.81E-224 | -0.5541 | -0.0117 (0.0003) | 1.16E-252 |
| EFNA2 | Ephrin-A2 | -0.0772 | -0.0278 (0.0049) | 1.69E-08 | -0.5312 | -0.0091 (0.0003) | 1.35E-216 | -0.5529 | -0.0084 (0.0002) | 3.15E-251 |
| CRABP2 | Cellular retinoic acid-binding protein 2 | -0.081 | -0.0414 (0.0074) | 2.21E-08 | -0.4069 | -0.0112 (0.0004) | 8.88E-138 | -0.4583 | -0.0114 (0.0004) | 1.33E-193 |
| HSPB6 | Heat shock protein beta-6 | -0.0692 | -0.041 (0.0073) | 2.35E-08 | -0.5076 | -0.0128 (0.0004) | 6.51E-186 | -0.4937 | -0.0109 (0.0004) | 2.83E-180 |
| VASN | Vasorin | -0.0888 | -0.0205 (0.0037) | 2.45E-08 | -0.3395 | -0.0046 (0.0002) | 2.03E-92 | -0.3392 | -0.0041 (0.0002) | 3.50E-98 |
| B4GALT1 | Beta-1,4-galactosyltransferase 1 | -0.0729 | -0.0214 (0.0038) | 2.73E-08 | -0.4407 | -0.0058 (0.0002) | 8.02E-138 | -0.4314 | -0.0051 (0.0002) | 9.76E-140 |
| GABARAP | Gamma-aminobutyric acid receptor-associated protein | -0.0947 | -0.0244 (0.0044) | 2.98E-08 | -0.4869 | -0.0075 (0.0002) | 1.44E-177 | -0.6207 | -0.0083 (0.0002) | 4.98E-315 |
| IL15RA | Interleukin-15 receptor subunit alpha | -0.073 | -0.0383 (0.0069) | 3.51E-08 | -0.4396 | -0.0102 (0.0004) | 2.04E-129 | -0.52 | -0.011 (0.0003) | 3.42E-207 |
| EPHB4 | Ephrin type-B receptor 4 | -0.072 | -0.0302 (0.0055) | 4.62E-08 | -0.4415 | -0.0084 (0.0003) | 4.51E-140 | -0.4348 | -0.0074 (0.0003) | 1.18E-144 |
| UNC5B | Netrin receptor UNC5B | -0.0794 | -0.0301 (0.0056) | 7.48E-08 | -0.4751 | -0.0091 (0.0003) | 3.90E-159 | -0.5033 | -0.0085 (0.0003) | 5.09E-189 |
| AMBP | Alpha-1-microglobulin | -0.084 | -0.0224 (0.0042) | 7.86E-08 | -0.4258 | -0.0063 (0.0002) | 3.53E-136 | -0.4805 | -0.0063 (0.0002) | 1.43E-189 |
| TAGLN | Transgelin | -0.0633 | -0.0354 (0.0066) | 7.98E-08 | -0.5736 | -0.0126 (0.0004) | 2.15E-230 | -0.5601 | -0.0105 (0.0003) | 3.27E-210 |
| AIF1L | Allograft inflammatory factor 1-like | -0.0669 | -0.0273 (0.0051) | 8.28E-08 | -0.4596 | -0.0078 (0.0003) | 1.84E-142 | -0.4507 | -0.0068 (0.0003) | 2.82E-140 |
| XXYLT1 | Xyloside xylosyltransferase 1 | -0.0748 | -0.0267 (0.005) | 8.78E-08 | -0.341 | -0.0058 (0.0003) | 8.09E-78 | -0.3364 | -0.0051 (0.0003) | 2.00E-79 |
| SERPINF1 | Pigment epithelium-derived factor | -0.0695 | -0.0176 (0.0033) | 1.36E-07 | -0.3319 | -0.0043 (0.0002) | 3.14E-95 | -0.3629 | -0.0042 (0.0002) | 2.14E-126 |
| MAP2K2 | Dual specificity mitogen-activated protein kinase kinase 2 | -0.084 | -0.0445 (0.0084) | 1.41E-07 | -0.3862 | -0.0117 (0.0005) | 5.85E-113 | -0.3532 | -0.0089 (0.0004) | 9.25E-87 |
| CAPG | Macrophage-capping protein | -0.0774 | -0.0452 (0.0086) | 1.46E-07 | -0.4178 | -0.0114 (0.0005) | 2.10E-103 | -0.4728 | -0.0113 (0.0004) | 3.13E-137 |
| EPHA1 | Ephrin type-A receptor 1 | -0.0728 | -0.0445 (0.0085) | 1.99E-07 | -0.283 | -0.0088 (0.0005) | 5.36E-61 | -0.3036 | -0.0085 (0.0004) | 1.19E-75 |
| IGFLR1 | IGF-like family receptor 1 | -0.0722 | -0.0455 (0.0088) | 2.44E-07 | -0.4744 | -0.0143 (0.0005) | 5.14E-160 | -0.5687 | -0.0153 (0.0004) | 9.54E-259 |
| EFEMP1 | EGF-containing fibulin-like extracellular matrix protein 1 | -0.0712 | -0.0243 (0.0047) | 2.53E-07 | -0.4354 | -0.0063 (0.0003) | 2.16E-106 | -0.5645 | -0.0075 (0.0002) | 6.01E-209 |
| DLK2 | Protein delta homolog 2 | -0.0682 | -0.0356 (0.0069) | 2.69E-07 | -0.4989 | -0.012 (0.0004) | 9.44E-185 | -0.4489 | -0.0094 (0.0003) | 1.84E-148 |
| IGFBP4 | Insulin-like growth factor-binding protein 4 | -0.0735 | -0.0249 (0.0049) | 3.33E-07 | -0.3655 | -0.0066 (0.0003) | 3.38E-109 | -0.4215 | -0.0067 (0.0002) | 1.18E-151 |
| VIT | Vitrin | -0.0664 | -0.0302 (0.0059) | 3.91E-07 | -0.3833 | -0.008 (0.0004) | 2.63E-105 | -0.3584 | -0.0067 (0.0003) | 1.30E-98 |
| PENK | Proenkephalin-A | -0.0753 | -0.0432 (0.0085) | 4.12E-07 | -0.3396 | -0.0091 (0.0005) | 1.13E-66 | -0.3779 | -0.0089 (0.0004) | 5.96E-85 |
| LRP10 | Low-density lipoprotein receptor-related protein 10 | -0.0682 | -0.0225 (0.0045) | 4.76E-07 | -0.4505 | -0.0069 (0.0003) | 5.60E-142 | -0.429 | -0.0058 (0.0002) | 1.05E-133 |
| CD93 | Complement component C1q receptor | -0.0684 | -0.0236 (0.0047) | 5.91E-07 | -0.2957 | -0.0051 (0.0003) | 4.17E-67 | -0.2939 | -0.0047 (0.0002) | 8.05E-76 |
| IL18BP | Interleukin-18-binding protein | -0.0645 | -0.029 (0.0058) | 6.38E-07 | -0.3747 | -0.0072 (0.0003) | 1.79E-88 | -0.4523 | -0.008 (0.0003) | 6.75E-150 |
| NPDC1 | Neural proliferation differentiation and control protein 1 | -0.0654 | -0.0295 (0.0059) | 6.87E-07 | -0.4681 | -0.0097 (0.0003) | 1.42E-162 | -0.4648 | -0.0083 (0.0003) | 1.17E-158 |
| WFDC2 | WAP four-disulfide core domain protein 2 | -0.0572 | -0.0349 (0.007) | 7.11E-07 | -0.5846 | -0.0141 (0.0004) | 1.56E-260 | -0.6098 | -0.0131 (0.0003) | 2.05E-304 |
| VWC2 | Brorin | -0.0691 | -0.027 (0.0055) | 8.57E-07 | -0.4736 | -0.0088 (0.0003) | 1.70E-157 | -0.4511 | -0.0072 (0.0003) | 3.25E-139 |
| RARRES2 | Retinoic acid receptor responder protein 2 | -0.0704 | -0.0284 (0.0058) | 9.13E-07 | -0.4542 | -0.0096 (0.0003) | 3.38E-167 | -0.5357 | -0.01 (0.0003) | 4.81E-252 |
| RNASE6 | Ribonuclease K6 | -0.0647 | -0.0459 (0.0093) | 9.40E-07 | -0.4655 | -0.0148 (0.0005) | 4.54E-152 | -0.5481 | -0.0156 (0.0004) | 2.94E-236 |
| PI3 | Elafin | -0.0566 | -0.0496 (0.0102) | 1.17E-06 | -0.4154 | -0.0155 (0.0006) | 1.64E-138 | -0.4343 | -0.0148 (0.0005) | 5.46E-171 |
| CALCOCO2 | Calcium-binding and coiled-coil domain-containing protein 2 | -0.069 | -0.0224 (0.0046) | 1.28E-06 | -0.3562 | -0.006 (0.0003) | 4.21E-98 | -0.314 | -0.0047 (0.0002) | 2.57E-80 |
| SPINK7 | Serine protease inhibitor Kazal-type 7 | -0.0628 | -0.0351 (0.0072) | 1.29E-06 | -0.4028 | -0.0105 (0.0004) | 4.92E-125 | -0.2811 | -0.0064 (0.0004) | 1.35E-60 |
| MIA | Melanoma-derived growth regulatory protein | -0.0641 | -0.0246 (0.0052) | 2.28E-06 | -0.2978 | -0.0054 (0.0003) | 1.65E-61 | -0.1855 | -0.0029 (0.0003) | 2.70E-24 |
| ASGR1 | Asialoglycoprotein receptor 1 | -0.0629 | -0.0284 (0.0061) | 2.79E-06 | -0.4406 | -0.0099 (0.0003) | 1.62E-160 | -0.4976 | -0.01 (0.0003) | 2.31E-227 |
| CLMP | CXADR-like membrane protein | -0.0587 | -0.034 (0.0073) | 2.80E-06 | -0.4428 | -0.0116 (0.0004) | 1.82E-155 | -0.4516 | -0.0104 (0.0004) | 1.13E-165 |
| CPLX2 | Complexin-2 | -0.0687 | -0.0356 (0.0076) | 3.05E-06 | -0.511 | -0.0135 (0.0004) | 1.62E-192 | -0.4491 | -0.0096 (0.0004) | 4.77E-125 |
| ATOX1 | Copper transport protein ATOX1 | -0.0954 | -0.0389 (0.0083) | 3.15E-06 | -0.3392 | -0.0104 (0.0005) | 2.25E-91 | -0.4176 | -0.0106 (0.0004) | 2.52E-128 |
| FABP3 | Fatty acid-binding protein, heart | -0.0529 | -0.0435 (0.0094) | 3.93E-06 | -0.4051 | -0.0157 (0.0005) | 1.78E-170 | -0.4974 | -0.0166 (0.0004) | 2.90E-268 |
| SMOC1 | SPARC-related modular calcium-binding protein 1 | -0.0671 | -0.0238 (0.0052) | 4.60E-06 | -0.4404 | -0.0074 (0.0003) | 2.96E-120 | -0.4244 | -0.0061 (0.0003) | 1.95E-109 |
| TMPO | Lamina-associated polypeptide 2, isoforms beta/gamma | -0.0768 | -0.039 (0.0085) | 5.20E-06 | -0.3591 | -0.0108 (0.0005) | 4.41E-94 | -0.4851 | -0.0127 (0.0004) | 5.88E-179 |
| DDOST | Dolichyl-diphosphooligosaccharide--protein glycosyltransferase 48 kDa subunit | -0.0691 | -0.025 (0.0055) | 5.34E-06 | -0.383 | -0.0072 (0.0003) | 5.18E-102 | -0.3842 | -0.0064 (0.0003) | 1.92E-106 |
| CST2 | Cystatin-SA | -0.0666 | -0.0537 (0.0119) | 6.29E-06 | -0.2963 | -0.0122 (0.0007) | 8.93E-61 | -0.1812 | -0.0062 (0.0006) | 1.24E-21 |
| NRXN3 | Neurexin-3-beta | -0.0649 | -0.0265 (0.0059) | 6.71E-06 | -0.3296 | -0.0064 (0.0004) | 2.00E-69 | -0.277 | -0.0045 (0.0003) | 8.94E-46 |
| SMOC2 | SPARC-related modular calcium-binding protein 2 | -0.0568 | -0.0272 (0.0061) | 7.97E-06 | -0.4313 | -0.0088 (0.0004) | 9.48E-126 | -0.3614 | -0.0063 (0.0003) | 3.47E-82 |
| NEGR1 | Neuronal growth regulator 1 | -0.0533 | -0.0172 (0.0039) | 8.57E-06 | -0.2982 | -0.0038 (0.0002) | 3.86E-56 | -0.1808 | -0.0018 (0.0002) | 1.09E-17 |
| Negative significant correlations, median^f^ | | -0.0855 | | | -0.4668 | | | -0.4697 | | |

^a^ Proteins were identified through linear regression of LDPred PRS for kidney function on 4,877 proteins measured at visit 3 and visit 5 with adjusting for age at the corresponded visits, sex, center, and first 10 genetic principal components. A total of 108 proteins were significantly associated with LDPred PRS at both visit 3 and visit 5. The threshold of significance was Bonferroni corrected: p = 1.02 × 10^-5^. Visit 3 was conducted during 1993-1995 when the mean age of study population was 60.4 years and visit 5 was conducted during 2011-2013 when the mean age of study population was 75.9 years.

^b^ LDPred PRS was constructed using LDPred algorithm, a Bayesian approach utilizes GWAS summary statistics to compute the posterior mean effect sizes for the genetic variants by assuming a prior of the joint effect sizes and incorporating the LD structure of the reference population.

^c^ Linear regression of LDPred PRS, eGFRcr measured at visit 3, and eGFRcys measured at visit 3 on proteins measured at visit 3, with adjusting for age at visit 3, sex, center, and first 10 genetic principal components.

^d^ Linear regression of LDPred PRS, eGFRcr measured at visit 5, and eGFRcys measured at visit 5 on proteins measured at visit 5, with adjusting for age at visit 5, sex, center, and first 10 genetic principal components.

^e^ P values of Wilcoxon signed rank test for the comparison between correlations of proteins with LDPred PRS and with eGFR were 2.58E-19 for eGFRcr and 1.15E-18 for eGFRcys at visit 3, and were 3.90E-19 for eGFRcr and 6.99E-19 for eGFRcys at visit 5.

^f^ P values of Wilcoxon signed rank test for the comparison between visit 3 and visit 5 correlations of proteins with LDPred PRS was 1.03E-13, and that of proteins with eGFR were 8.71E-19 for eGFRcr and 2.13E-18 for eGFRcys.

eGFRcr: estimated glomerular filtration rate based on creatinine; eGFRcys: estimated glomerular filtration rate based on cystatin C; PRS: polygenic risk score
